# miR-573 rescues endothelial dysfunction during dengue infection under PPARγ regulation

**DOI:** 10.1101/2021.08.30.458308

**Authors:** Shefali Banerjee, Kong Hao Yuin, Chin Wei Xin, Justin Jang Hann Chu

## Abstract

Early prognosis of abnormal vasculopathy is essential for effective clinical management of severe dengue patients. An exaggerated interferon (IFN) response and release of vasoactive factors from endothelial cells cause vasculopathy. This study shows that dengue 2 (DENV2) infection of human umbilical vein endothelial cells (HUVEC) results in differentially regulated miRNAs important for endothelial function. miR-573 was significantly down-regulated in DENV2-infected HUVEC due to decreased Peroxisome Proliferator Activator Receptor Gamma (PPARγ) activity. Restoring miR-573 expression decreased endothelial permeability by suppressing the expression of vasoactive angiopoietin 2 (ANGPT2). We also found that miR-573 suppressed the proinflammatory IFN response through direct downregulation of toll like receptor 2 (TLR2) expression. Our study provides a novel insight into miR-573 mediated regulation of endothelial function during DENV2 infection which can be further translated into a potential therapeutic and prognostic agent for severe dengue patients.

---

Dengue is an escalating public health concern with 100-200 million symptomatic infections occurring annually across the globe [1]. Dengue virus (DENV) belongs to *Flaviviridae* family and has four distinct serotypes which are transmitted by the *Aedes* mosquitoes. The disease is usually subclinical or presented with symptoms such as fever, headache, myalgia and arthralgia in most patients. However, certain patients develop a severe and potentially fatal form of the disease called as dengue shock syndrome due to vasculopathy. Increasing trends in urbanisation and international travel have contributed in global distribution of the different serotypes, resulting in hyperendemicity and increasing the risk of severe disease which is more likely during secondary infections by different DENV serotypes [2]. Severe dengue is associated with hemodynamic imbalance resulting in excessive plasma leakage in pleural cavities and hypovolemic shock [3]. Predicting the onset of severe dengue is clinically challenging resulting in unnecessary hospitalization and over burdening of the public healthcare systems, especially in the endemic regions. The economic burden of the disease is estimated to be US$ 3-4 billion in these regions highlighting the need for better preventive and diagnostic approaches for efficient clinical management of this disease [4, 5]. The current laboratory and clinical markers for predicting the occurrence of severe dengue in patients have paired poorly in clinical settings [6, 7]. Designing better diagnostic approaches will require a detailed understanding of the mechanisms contributing to the pathophysiology of the disease.

During severe dengue, antibody dependent enhancement (ADE) [8-10], abnormal T cell function [11], platelet activation [12] and mast cell degranulation induce vasoactive mediators such as, TNFα, VEGF, IP-10, GM-CSF, IL-1α, IL-1β and IL-8, CXCL1, HMGB1, MCP1, metalloproteinases which disrupt the endothelial barrier integrity leading to vascular leakage [13]. DENV non-structural protein 1 (NS1) also induces endothelial permeability by directly disrupting the endothelial glycocalyx layer [14]. Circulating levels of endothelial activation markers like sICAM1, sVCAM1, claudin5, as well as the components of the endothelial glycocalyx layer (syndecan, chondroitin sulphate) are elevated in severe dengue patients suggesting the involvement of a dysregulated endothelial barrier function [15-19]. Severe dengue patients present early signs of endothelial dysfunction as seen by the elevated Reactive Hyperemia Index (RHI) which was associated with a 4-fold risk of severe dengue [20]. Identifying such factors directly associated with endothelial function during severe dengue is essential and requires focused exploration of endothelial cell function during dengue infection.

MicroRNAs (miRNA) are a class of small RNAs which have recently been found to be involved in regulating endothelial cell (EC) function [21]. miRNAs bind to the 3’UTR of target mRNAs and post transcriptionally regulate protein production [22]. miR-125, miR-155 and miR221/222 are some of the miRNAs which are involved in regulating key EC functions like angiogenesis, inflammation and permeability [23]. Recent studies have also highlighted the role of miRNAs during dengue infection in both *in vitro* and clinical studies [24-27]. However, these studies focused on miRNA expression in cells of non-EC origin and the role of endothelial miRNAs during DENV infection is yet to be explored. Proinflammatory stimuli which regulate EC miRNA expression, especially during vascular pathology, are also common in severe dengue infection [28]. Identifying EC miRNAs which regulate pathways associated with inflammation and permeability responses may provide common regulatory networks between the multiple factors mediating EC permeability.

In this study we show that DENV2 infection of HUVEC cells resulted in differential regulation of miRNAs involved in EC function. HUVEC has been widely used for vascular pathology studies and is a permissive host for DENV infection [29, 30]. Several of the differentially regulated miRNAs were required for maintaining vascular tone and endothelial cell homeostasis. We further elaborated on the role of miR-573 in regulation of endothelial cell function during DENV2 infection. We also provide some preliminary evidence of regulation of miR-573 expression through Peroxisome Proliferator Activator Receptor Gamma (PPARγ) at the transcriptional level.

## Results

### DENV2 infection of HUVEC modulates endothelial miRNA expression

Small RNA sequencing was carried out to identify the changes in the miRNA expression profile in HUVEC cells upon DENV2 infection (MOI 10). HUVEC cells have been widely used as an in vitro model for vascular studies and are susceptible to DENV infection [29, 30]. We identified 188 miRNAs to be differentially regulated during DENV2-infected HUVEC (Figure 1a). 38 miRNAs from our screen have previously been reported to be differentially regulated during DENV infection of other non-endothelial cell types indicating the robustness of our screen (Figure 1 – figure supplement 1). A complete list of the differentially expressed miRNAs at 24hpi and 48hpi are tabulated in (Supplementary File 1). We selected a few of the most highly significant and differentially expressed miRNAs for qRT-PCR analysis and we observed a similar trend in their expression profiles further validating the robustness of the sequencing analysis (Figure 1 – figure supplement 1). Gene ontology analysis of the differentially regulated miRNAs using Diana miRpath tool [31] identified pathways associated with immune response (JAK-STAT, cytokine and chemokine signaling), metabolism (Wnt, mTOR, Fatty acid synthesis, PI3-AKT signaling), p53 signaling and endothelial permeability response (actin cytoskeleton remodeling, gap and tight junctions, adherens junction and ECM-receptor signaling) (Figure 1 – figure supplement 1). To further complement the RNA-seq data we carried out a mRNA microarray to determine the overall changes in global mRNA expression levels. (Figure 1b). A complete list of the differentially expressed mRNAs are tabulated in (Supplementary File 2). We superimposed the miRNA and mRNA datasets and identified several putative miRNA-mRNA associations which exhibited a reciprocal relationship in their expression levels. We also identified miRNAs like miR-15, miR-26, miR-30, miR-365, and miR-483 which regulated mRNAs involved in pathways like gap and tight junctions, adherens junction and extracellular matrix receptor (ECM-receptor) signaling (Figure 1c) in endothelial cells [32-35]. We observed that miR-15 (Figure 1d), miR-26 (Figure 1e), miR-30 (Figure 1f), miR-365 (Figure 1g), and miR-483 (Figure 1h) were down-regulated in HUVEC at 24 hpi and some of their putative mRNA targets were up-regulated in our microarray dataset (Supplementary File 3) further emphasizing on the reciprocal relationship between miRNA-mRNA expressions. These observations indicate that DENV2 infection results in an altered miRNA expression profile in endothelial cells which may be responsible in modulating the endothelial cell permeability and activation responses during DENV infection. We also identified miR-573 to be significantly down-regulated at 24hpi which had putative mRNA targets involved in endothelial permeability and activation responses. Previous reports have shown that miR-573 mitigates the proinflammatory response in a rheumatoid arthritis disease model and has an endothelial protective function [36]. We found miR-573 to be downregulated in HUVEC cells throughout the course of DENV2 infection (Figure 2a). In order to eliminate any cell line biases we also studied the expression of miR-573 during DENV2 infection in HMEC-1 cell line, another well-established in vitro endothelial cell model. As in DENV2-infected HUVEC, miR-573 expression was also downregulated in HMEC-1 cells at 48hr and 72hr post infection (Figure 1 – figure supplement 1).

**Figure 1:**
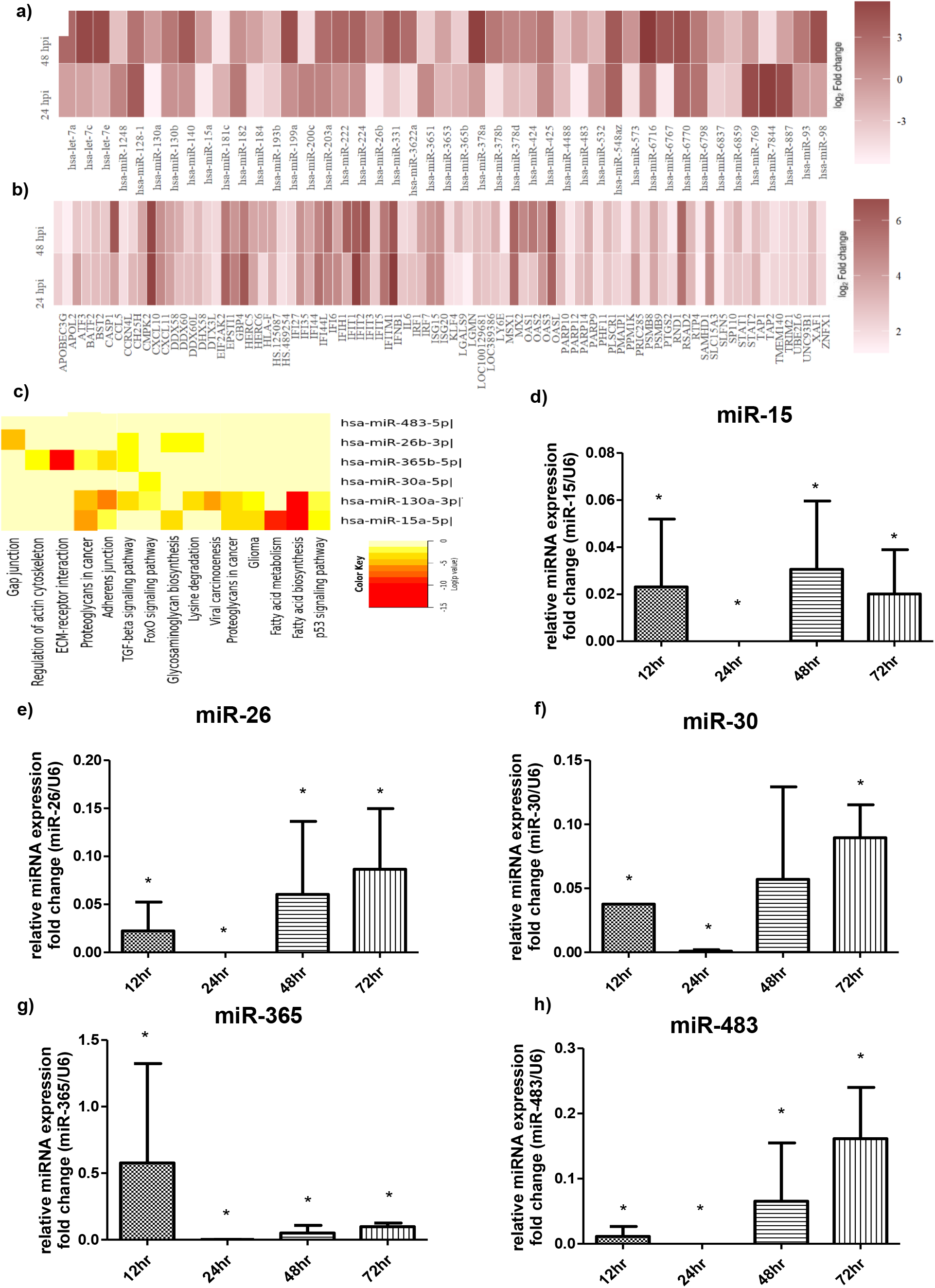
DENV2 infection alters the miRNA expression in HUVEC cells. **a)** Heat map representing the differentially expressed miRNAs in DENV2-infected HUVEC at 24 and 48 hpi. **b)** A heat map depicting the overall change in the mRNA expression in response to DENV2-infection. **c)** DIANA mirPath pathway enrichment analysis (Fisher’s exact test (p < 0.05) and False discovery rate (FDR=0.01)) identified potential pathways regulating endothelial function like gap junction signaling, tight junction signaling, fatty acid synthesis etc. qRT-PCR analysis of miRNA expression levels for miR-15 **(d)**, miR-26 **(e)**, miR-30 **(f)**, miR-130 **(g)**, miR-365, and **(h)** miR-483 during DENV2-infected cells with respect to mock-infected control. Data represented mean ± SEM and are obtained from three individual biological replicates. Student’s t test was carried out between individual groups to determine the statistical difference. *\*P<0*.*05*

**Figure 2:**
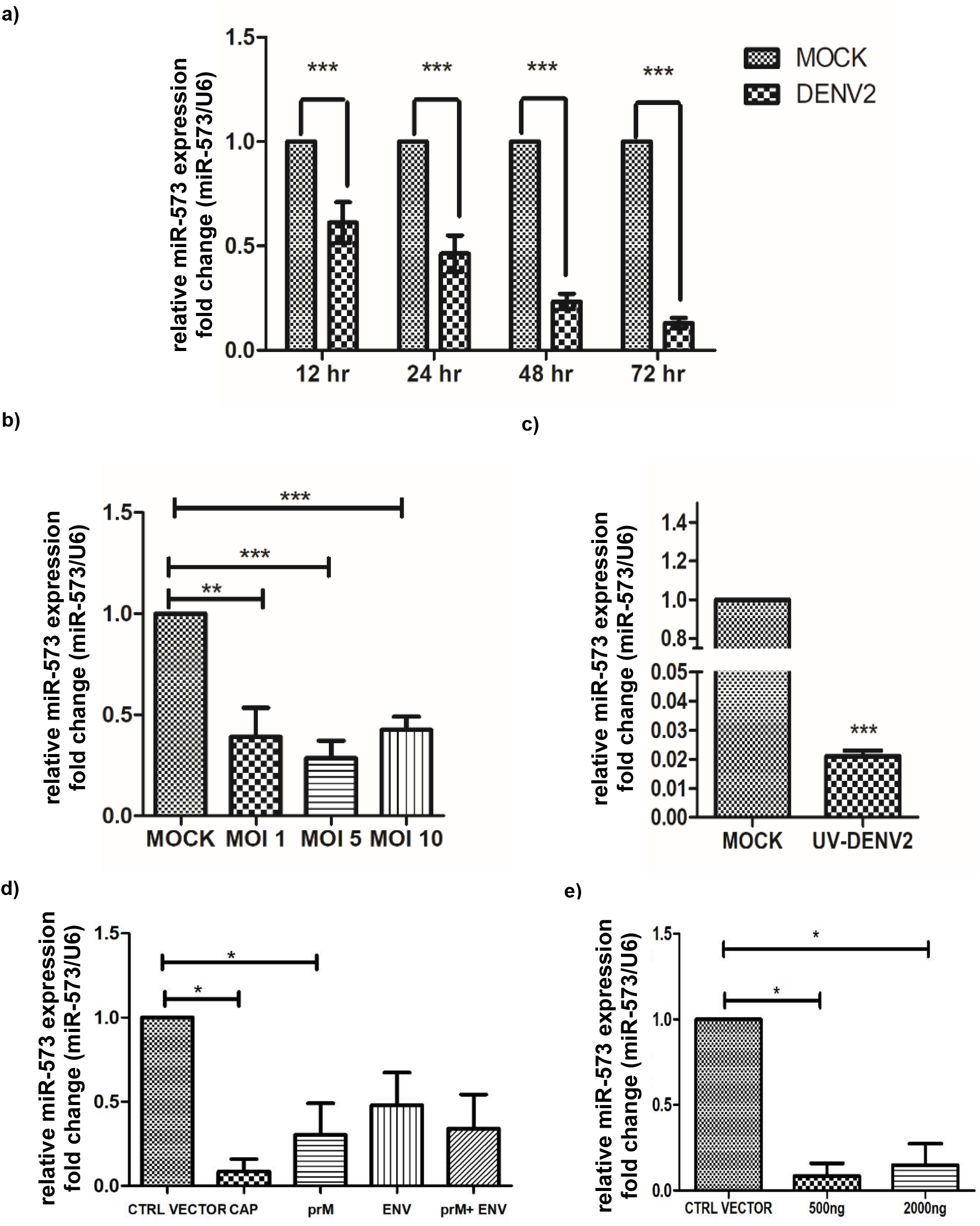
Regulation of miR-573 expression does not directly depend on active viral replication. qRT-PCR analysis of miR-573 transcript levels in cells when infected with DENV2 **a)** at different time intervals post infection and **b)** at different MOIs relative to mock-infected cells. **c)** HUVEC cells were challenged with UV-inactivated DENV2 virus and miR-573 expression levels were measured relative to mock-infected cells. **d)** miR-573 expression in HUVEC cells transiently transfected with recombinant DENV2 viral proteins relative to cells transfected with the control vector Data represented mean ± SEM and are obtained from three individual biological replicates. Student’s t test was carried out between individual groups to determine the statistical difference. *\*P<0*.*05, **P<0*.*01, ***P,0*.*001*. ENV-envelope, cap-capsid, prM-precursor membrane

### Regulation of miR-573 expression is independent of DENV2 viral replication

We observed that miR-573 was downregulated in DENV2-infected HUVEC at different multiplicities of infection (MOI 1, 5 & 10) relative to the mock-infected cells (Figure 2b). However, we did not observe a significant difference in miR-573 expression between the different MOIs. To further determine whether miR-573 expression is dependent on active viral replication, HUVEC cells were infected with an UV-inactivated DENV2 virus. miR-573 expression was reduced in cells treated with the UV-inactivated virus (Figure 2c). These observations probably suggest that dysregulation of miR-573 expression in DENV2-infected HUVEC is independent of active viral replication. Downregulation of miR-573 expression in response to the replication defective UV-inactivated virus lead us to hypothesize whether its expression is modulated by the viral proteins which remain intact upon UV exposure. We transiently transfected HUVEC cells with the DENV2 structural gene constructs (envelope, prM and capsid proteins) and measured the change in miR-573 expression (Figure 2 – figure supplement 1). We observed a decrease in miR-573 expression for all the three structural proteins and transfection with the capsid construct resulted in greater downregulation (Figure 2d) compared to the other constructs. However, we did not observe a dose dependent decrease in miR-573 expression in response to the increasing concentrations of the capsid protein (Figure 2e) suggesting that regulation of miR-573 expression might depend on additional factors in addition to this protein.

### miR-573 expression is under the transcriptional control of Peroxisome Proliferator Activator Receptor Gamma (PPARγ)

miR-573 is an intergenic miRNA located between DHX15 and PPARγCA1 genes on chromosome 4 (Figure 3 –figure supplement 1). We also identified several markers of transcriptionally active regions like the histone modification signals (140 H3K4me3 & H3K27Ac) and DNase1 hypersensitive regions located upstream of miR-573 coding region using the miRStart computational tool (Figure 3 – figure supplement 1) [37]. We also identified Peroxisome Proliferator Activator Receptor Gamma (PPARγ) transcription factor binding sites located within 2kb upstream of miR-573 coding region (Figure 3 – figure supplement 1). PPARγ which acts as a nuclear hormone receptor has been shown to suppress the proinflammatory response in endothelial cells in a sepsis disease model [38]. We observed a significant reduction in PPARγ protein levels in DENV2-infected HUVEC (Figure 3a & 3b). PPARγ is a ligand activated transcriptional factor which upon activation undergoes nuclear translocation and binds to the PPAR response elements (PRE) upstream of its target genes. PPARγ levels in the nuclear fraction of DENV2-infected HUVEC was significantly lower than mock-infected cells (Figure 3c). To further determine whether modulation of PPARγ activity influences miR-573 transcript levels, HUVEC cells were treated with varying concentrations of rosiglitazone, which is a PPARγ agonist. A dose dependent increase in miR-573 expression levels was observed upon rosiglitazone treatment (Figure 3d). We also observed that treating DENV2-infected cells with rosiglitazone pre and post infection resulted in an increase in miR-573 expression as opposed to the solvent treated cells (Figure 3 – figure supplement 2). Whereas inhibiting PPARγ activity with antagonist GW9662 resulted in a dose dependent decrease in miR-573 expression in HUVEC cells (Figure 3e). These observations suggest that miR-573 expression depends on PPARγ activity. Transient overexpression of PPARγ resulted in a significant increase in miR-573 expression in DENV2-infected HUVEC (Figure 3f & 3g). The PPARγ binding site was predicted to be 1536 bp upstream of miR-573 primary transcript, located between 2452178 and 24521826 bp on chromosome 4. We carried out a chromatin immunoprecipitation assay to show direct binding of PPARγ to the predicted site upstream of the miR-573 coding region (Figure 3h).

**Figure 3:**
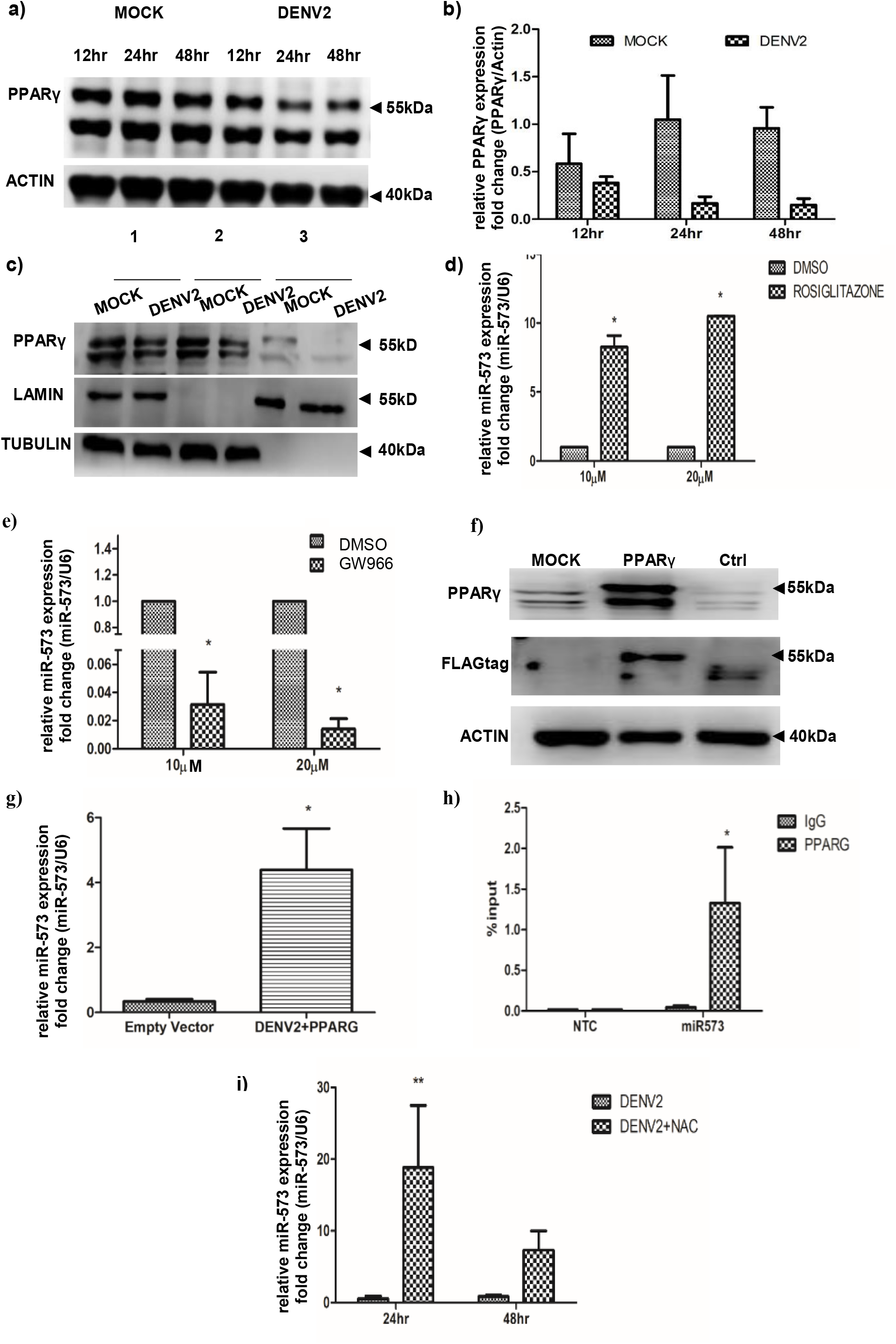
miR-573 expression is under the transcriptional control of PPARγ. **a)** Western blot analysis of PPARγ expression upon DENV2-infected HUVEC at 12,24 and 48 hpi. **b)** Histogram representing the densitometric analysis of the western blot for PPARγ expression. **c)** Cytoplasmic (lane 2) and nuclear fractions (lane 3) of DENV2 and mock-infected cells were analyzed by western blot analysis for PPARγ translocation. Whole cell lysates are represented in lane 1 of the western blot for mock and DENV2-infected cells. HUVEC cells were treated with different concentrations of **d)** rosiglitazone and **e)** GW9662 and miR-573 expression was measured relative to solvent treated cells by qRT-PCR. **f)** Western blot comparing the PPARγ expression in transiently overexpressing, control and untreated cells. **g)** miR-573 expression levels in DENV2-infected HUVEC overexpressing PPARγ relative to infected cells transfected with the empty vector. **h)** CHIP assay to determine the PPARγ binding site for miR-573 on chromosome 4. Fold enrichment was determined relative to position P1 which had no PPARγ binding site. **i)** miR-573 expression levels upon NAC treatment followed by DENV2 infection. Data represented mean ± SEM and are obtained from three individual biological replicates.. Student’s t test was carried out between individual groups to determine the statistical difference. *\*P<0*.*05, **P<0*.*01*

### Oxidative stress induced during DENV2-infection of HUVEC regulates PPARγ activity

DENV2-infection resulted in a decrease in PPARγ expression levels and a parallel decrease in its nuclear translocation ability. This decrease in PPARγ activity correlated with the decrease in miR-573 expression levels. Treatment with PPARγ agonist rosiglitazone was able to increase PPARγ activity and a consequent increase in miR-573 expression. These observations suggest that DENV2-infection triggers a response which together modulates the abundance of endogenous PPARγ activating ligands as well as the PPARγ expression levels. Presence of Reactive Oxygen Species (ROS) has previously been reported to negatively regulate PPARγ expression in endothelial cells [38]. Oxidative stress response has also been linked to an indirect decrease in PPARγ activity due to the elimination of endogenous PPARγ ligands from the host cell [38]. DENV2 infection also induces an oxidative stress response in host cells [39] and it could be involved in the differential regulation of PPARγ and miR-573 expression in the infected cells. To determine whether oxidative stress plays a role in PPARγ expression levels, HUVEC cells were treated with N-acetyl cysteine (NAC) treatment to suppress ROS production (Figure 3 – figure supplement 2). NAC is an aminothiol and acts as a reducing agent as well as a synthetic precursor of intracellular cysteine and glutathione (GSH) antioxidant system [40]. We observed that upon NAC treatment PPARγ protein levels were elevated in DENV2-infected HUVEC (Figure 3 – figure supplement 2). Consequently, NAC treatment also resulted in an increase in miR-573 expression during DENV2 infection (Figure 3i). These findings suggest that regulation of PPARγ activity and its downstream target miR-573 depends on the oxidative stress response induced during DENV2 infection.

### Overexpression of miR-573 decreases DENV2 mediated endothelial permeability

Since miR-573 was down-regulated during DENV2 infection we further analyzed the effect of miR-573 overexpression on endothelial permeability upon DENV2 infection. HUVEC cells were transfected with LNA modified miR-573 mimic and a non-targeting mimic (NTC) was used as a control. miR-573 expression in mimic transfected cells was significantly higher than the control cells (Figure 4a). miR-573 overexpressing cells were infected with DENV2, following which the permeability characteristics of the endothelial monolayer were assayed. We observed an increase in transendothelial electrical resistance (TEER) in DENV2-infected endothelial monolayer overexpressing miR-573 (Figure 4b). Similarly, the dextran FITC influx across the infected monolayer was significantly lower in cells overexpressing miR-573 (Figure 4c). We also observed a similar effect in miR-573 overexpressing HMEC-1 cells upon DENV2 infection (Figure 4 – supplement 1). DENV2 infection of HUVEC results in disorganization of the intercellular adherens junction proteins like VE-cadherin [30, 41]. To determine whether miR-573 overexpression restores the VE-cadherin arrangement in DENV2 infected cells, VE-cadherin protein organization in the infected cells was visualized using immunofluorescence staining. Compared to the NTC transfected cells, the VE-cadherin proteins were more intact at the intercellular junctions upon miR-573 overexpression (Figure 4d & 4e). Inhibition of miR-573 activity by miR-573 LNA modified inhibitor resulted in no significant difference between the TEER measurements compared to NTC in DENV2-infected HUVEC. However, inhibition of miR-573 activity in mock treated HUVEC resulted in a significant difference in TEER values compared to the control (Figure 4 – figure supplement 1). No miR-573 binding sites were identified in the DENV2 genome miR-573 overexpression had no significant effect on viral protein expression as well as viral replication (Figure 4f & 4g). These observations suggest that the rescue in endothelial permeability upon miR-573 overexpression is not mediated through suppression of viral replication but rather through regulation of host factors which are induced upon viral infection.

**Figure 4:**
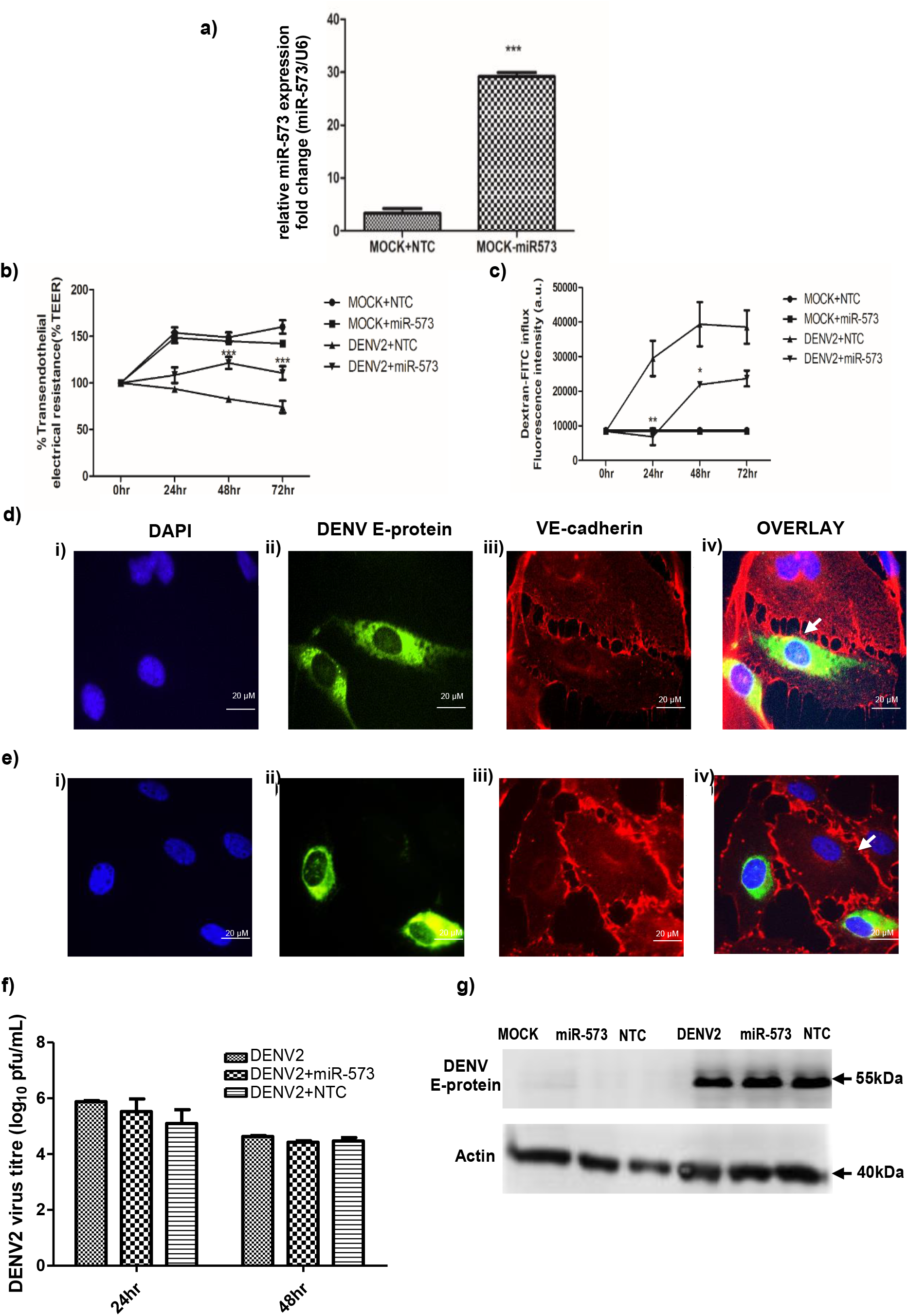
miR-573 reduces endothelial permeability in DENV2-infected HUVEC. **a)** HUVEC were transfected with miR-573 mimic (80nM) or the mimic non-targeting control (NTC, 80nM) and miR-573 expression levels were determined by qRT-PCR.HUVEC cells transfected with the mimic or the non-targeting control were seeded onto the transwell inserts and endothelial permeability was measured following mock or DENV2 infection. **b)** TEER measurements across DENV2-infected endothelial monolayer transfected with miR-573 mimic increased gradually from 26.3%, 35.2% and 28% at 24, 48 and 72hpi, respectively (n=3, * p<0.05, **p<0.01). **c)** Similarly, the dextran FITC influx across the DENV2-infected endothelial monolayer transfected with miR-573 mimic decreased from 73.2%, 21.3% and 36.6% at 24, 48 and 72hpi respectively (n=3, *p<0.05, **p<0.01). VE-cadherin (red) and DENV E-protein (green) were visualized using double immunofluorescence in DENV2-infected HUVEC. VE-cadherin junctions were disrupted and punctated in cells treated with the NTC **(d-iv)** as compared to the more distinct and regular arrangement observed in DENV2-infected HUVEC treated with **(e-iv)** miR-573 mimic **f)** DENV2 viral titers in DENV2-infected HUVEC treated with miR-573 mimic or the non-targeting mimic control were measured using plaque assay. No significant difference in viral titers was observed in the individual treatment groups. **g)** Western blot analysis of E-protein expression in DENV2-infected HUVEC treated with the mimic or the NTC showed no significant change. Data represented mean ± SEM and are obtained from three individual biological replicates. Student’s t test was carried out to determine the statistical difference. *\*P<0*.*05, **P<0*.*01, ***P<0*.*001*.

### miR-573 directly represses ANGPT2 expression and modulates endothelial permeability

ANGPT2 is one of the identified miR-573 targets obtained from three individual miRNA target prediction tools (Targetscan, Diana microT-CDS and miRDB) [42-44]. Earlier studies have reported that increase expression of ANGPT2 was responsible for increasing endothelial permeability during DENV2 infection [30, 45]. Our study also found that ANGPT2 expression levels were elevated in DENV2-infected HUVEC (Figure 6 – figure supplement 1). To determine whether miR-573 binds to the predicted miRNA recognition element (MRE) within the 3’UTR of ANGPT2, HUVEC cells were co-transfected with the mimic or the NTC and luciferase constructs carrying wild type or the mutant MRE sequence (Figure 5a). The luciferase expression in cells carrying the wild type MRE construct was significantly lower upon miR-573 overexpression with respect to the NTC (Figure 5b). No significant decrease in luciferase expression levels was observed from the luciferase construct carrying the mutant MRE between cells transfected with miR-573 and the NTC. Similarly, no significant difference was seen in luciferase expression upon inhibition of miR-573 activity for the individual ANGPT2-MRE constructs with respect to NTC (Figure 5 – figure supplement 1). ANGPT2 mRNA transcript levels were also downregulated upon miR-573 overexpression (Figure 5c). miR-573 overexpression resulted in a significant decrease in both intracellular and extracellular ANGPT2 protein levels upon DENV2 infection (Figure 5d-5f) whereas inhibition of miR-573 activity resulted in no significant difference in ANGPT2 protein expression levels (Figure 5 – figure supplement 1). To further validate whether the decrease in endothelial permeability observed upon miR-573 overexpression is mediated through suppression of ANGPT2 expression, recombinant ANGPT2 protein was exogenously added to the infected cells overexpressing miR-573. Addition of ANGPT2 resulted in a gradual decrease in TEER measurements in miR-573 overexpressing infected cells (Figure 5g). Similarly, the dextran FITC influx significantly increased in miR-573 overexpressing cells after the addition of ANGPT2 (Figure 5h). These observations suggest that miR-573 binds to the 3’UTR of ANGPT2 and suppresses its expression which may contribute to the decrease in endothelial permeability observed upon miR-573 overexpression.

**Figure 5:**
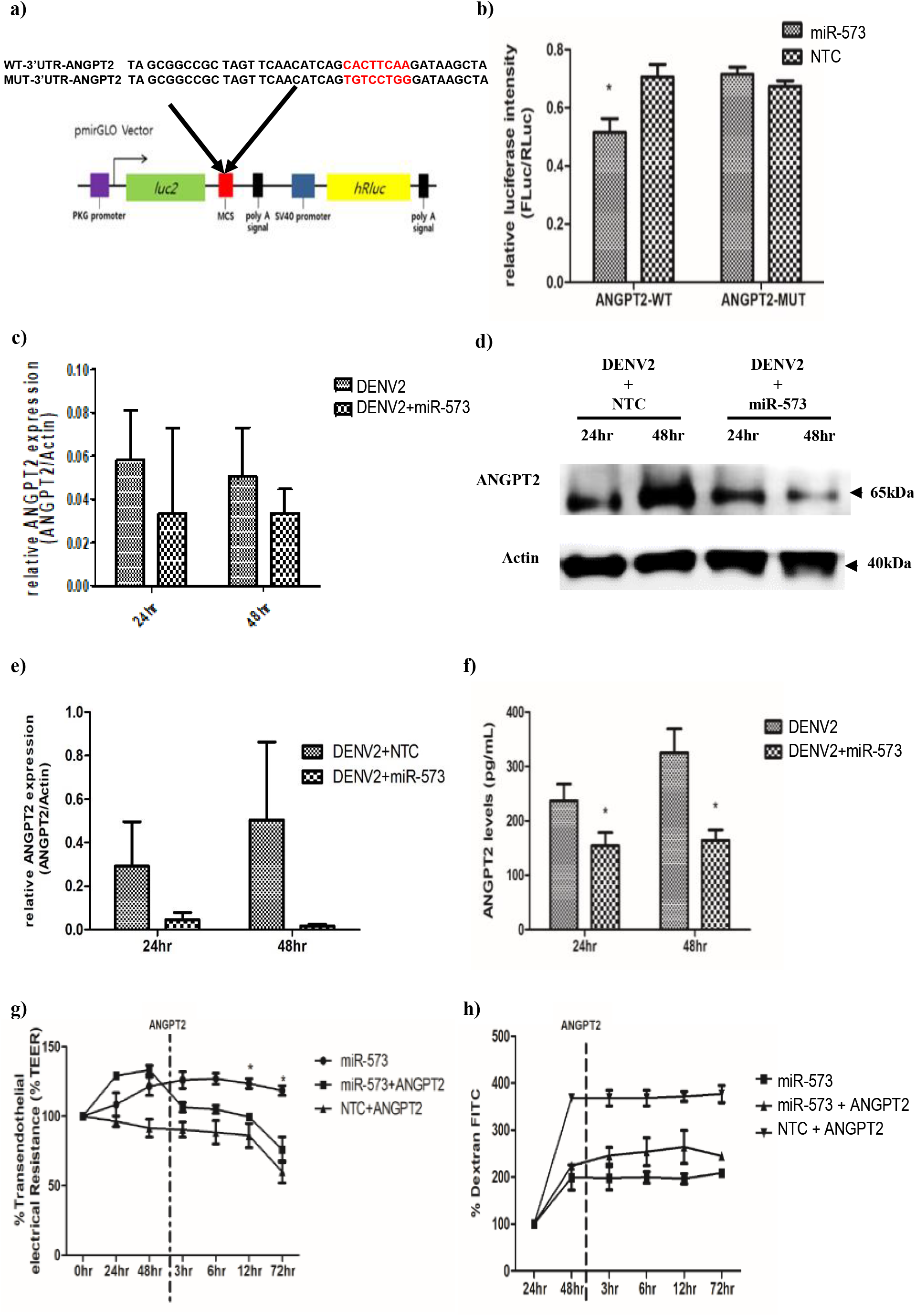
miR-573 targets ANGPT2 expression in DENV2-infected HUVEC. **a)** Diagrammatic representation of the WT-3’UTR-ANGPT2 and MUT-3’UTR-ANGPT2 constructs using the backbone of the pmiRGlO dual-luciferase miRNA target expression vector **b)** HUVEC were co-transfected with WT-3’UTR-ANGPT2 construct and miR-573 mimic or the non-targeting control (NTC) and the relative luciferase signal intensity (Fluc/Rluc) was measured at 24 hpt. HUVEC cells transfected with miR-573 mimic or the non-targeting control were infected with DENV2 at MOI 10 and cell lysates and supernatants were collected at regular time intervals for determining ANGPT2 mRNA and protein expression. **c)** A bar graph depicting the relative ANGPT2 mRNA transcript levels **d)** Western blot analysis of ANGPT2 protein levels upon miR-573 overexpression. **e)** A bar graph showing the densitometric analysis of the ANGPT2 protein expression. **f)** Bar graph showing the levels of extracellular ANGPT2 measured by ELISA. Recombinant ANGPT2 (30ng/ml) was added to the DENV2-infected HUVEC monolayer at 48 hpi and the **g)** TEER measurements and **h)** dextran FITC influx was measured at regular time intervals. Data represented mean ± SEM and are obtained from three individual biological replicates. Student’s t test was carried out between individual groups to determine the statistical difference. *\*P<0*.*05*.

### miR-573 regulates the proinflammatory response upon DENV2 infection of HUVEC

To further determine whether miR-573 affects the endothelial activation response in DENV2-infected HUVEC we measured the secreted levels of proinflammatory cytokines like IL-6, IL-1β and TNF-α from these cells. In miR-573 overexpressing DENV2-infected HUVEC cells IL-6 levels were significantly reduced as compared to NTC (Figure 6a). Whereas inhibition of miR-573 activity resulted in no significant decrease in IL-6 secretion from DENV2-infected HUVEC (Figure 5 – figure supplement 1). We did not observe a significant reduction in IL-1β and TNF-α levels between DENV2-infected HUVEC overexpressing miR-573 and the NTC (data not shown). The levels of these cytokines were below the detection limit of even in mock and DENV2-infected HUVEC in our experimental set up (data not shown). An exaggerated IFN response is responsible for increased endothelial permeability in various proinflammatory diseases [29, 46] and our target prediction analysis found miR-573 to target several of these IFN response genes (Supplementary File 3). RSAD2, OAS2, GBP1 etc., have been shown to be associated with an exaggerated endothelial activation response [47, 48]. The mRNA transcript levels of these IFN response genes were significantly lower in DENV2-infected cells overexpressing miR-573 with respect to NTC (Figure 6b). Endothelial activation promotes leukocyte transmigration across the monolayer and we further explored whether miR-573 overexpression reduces leukocyte transmigration across the infected monolayer. Human PBMC were co-incubated with HUVEC cells on transwell inserts and after 3hr of incubation, the percentage of transmigrating PBMCs in the lower chamber was determined by flow cytometry. Percentage of transmigrating CD14 and CD45 positive cells (CD14^+^ CD45^+^) across the DENV2-infected monolayer subjected to different treatments; DENV2 infection alone (Figure 6c), DENV2-infected with miR-573 mimic (Figure 6d) and DENV2-infected with NTC (Figure 6e) was evaluated. A significant decrease in the percentage of the transmigrating PBMC was observed across the DENV2-infected endothelium overexpressing miR-573 as compared to the NTC treated cells (Figure 6f). Inhibition of miR-573 activity resulted in no significant difference in the percentage of transmigrating PBMCs across the DENV2-infected monolayer with respect to NTC (Figure 5 – figure supplement 1). However, neither IL-6 nor the IFN response genes were validated targets of miR-573 suggesting that miR-573 might target factors responsible in IFN response regulation during DENV2 infection.

**Figure 6:**
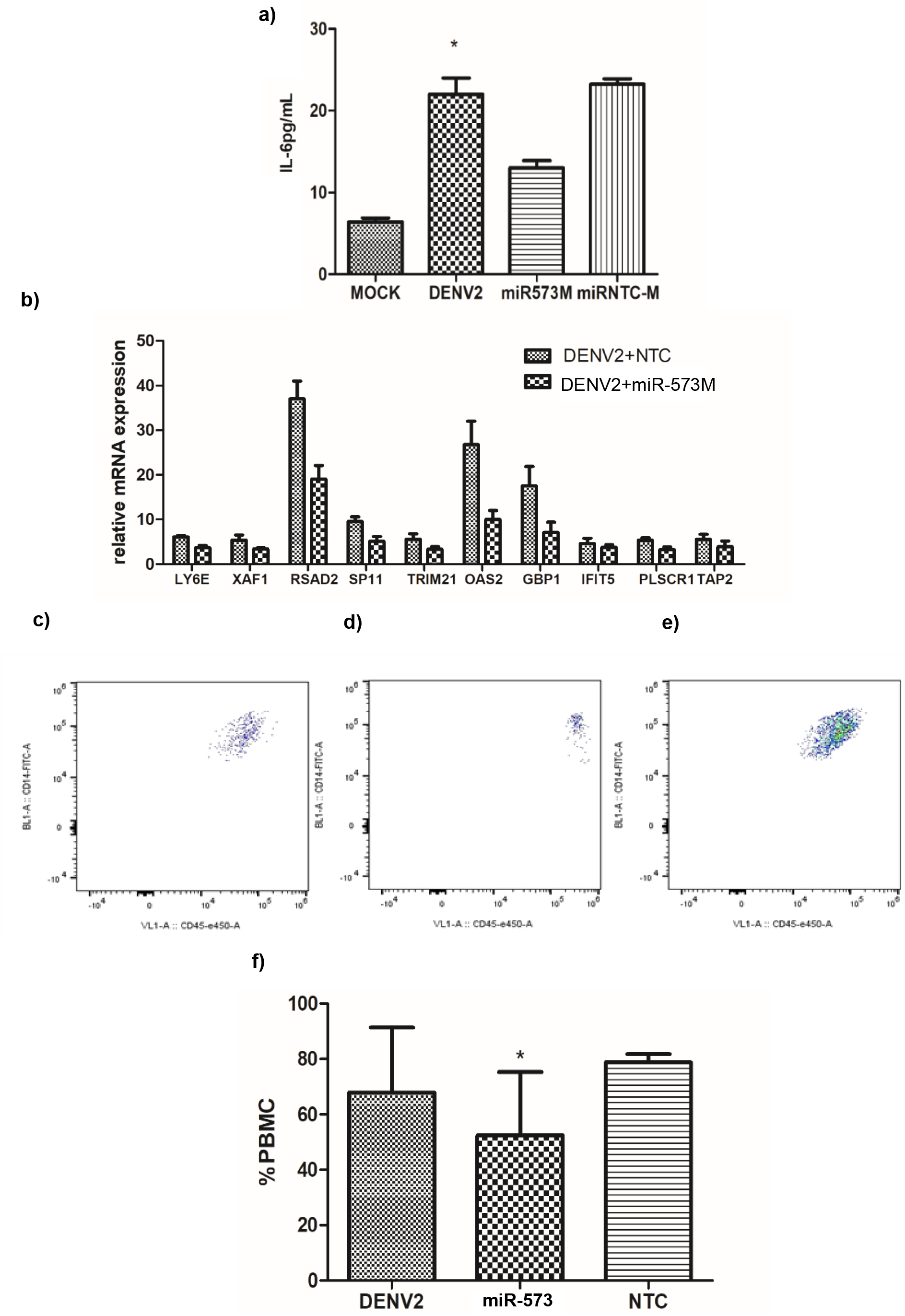
Overexpressing miR-573 suppresses the endothelial activation response. HUVEC cells were transfected with miR-573 mimic (80nM) or non-targeting mimic control (NTC, 80nM) prior to DENV2 infection (MOI 10). **a)** Secreted levels of IL-6 from DENV2-infected HUVEC were measured using ELISA at 24hpi. **b)** The mRNA transcript levels of endothelial activation markers were measured at 24hpi. Percentage of transmigrating leukocytes across the infected HUVEC monolayer upon **c)** DENV2 infection, **d)** NTC treatment and **e)** miR-573 mimic treatment was measured using flowcytometry. Representative images depicting the populations of CD14 and CD45 positive cells for each of the treatments are shown. **f)** A bar graph depicting the percentage of PBMC transmigrating across the infected HUVEC monolayer subjected to the different treatments. Data represented mean ± SEM and are obtained from three individual biological replicates. Student’s t test was applied to determine the statistical difference between individual groups *\*P<0*.*05*.

### miR-573 binds to the 3’UTR of TLR2 and modulates endothelial activation

Toll like receptors 2 and 4 (TLR2 and TLR4) were found to be the potential predicted targets of miR-573 which also induce IFN response upon activation. TLR2 signaling is responsible for DENV NS1 protein mediated pathogenesis in DENV [49]. We hypothesized that the observed decrease in the type 1 IFN response upon miR-573 overexpression may be a consequence of the post transcriptional repression of TLR2/4 expression. Both TLR2 and TLR4 mRNA and protein levels were elevated in DENV2-infected HUVEC (Figure 6 – figure supplement 1). To further validate whether miR-573 binds to the MRE present within the TLR2/4 3’UTRs we repeated the luciferase reporter assays with the TLR2/4-MRE constructs. The luciferase expression was significantly lower in HUVEC cells co-transfected with miR-573 and the luciferase construct carrying the wild type TLR2 MRE sequence (Figure 7a). Overexpressing miR-573 resulted in a progressive decline in TLR2 protein levels from 26.7% to 58.5% from 24 to 72hpi (Figure 7b & 7c). However, no significant decrease in luciferase expression was observed for the TLR4-MRE construct under similar conditions (Figure 7d). No change in TLR4 protein levels was observed in DENV2-infected HUVEC transfected with miR-573 mimic as compared to cells transfected with the NTC (Figure 7e & 7f). Therefore, miR-573 only binds to the 3’UTR of TLR2 and decreases its expression and not to the predicted MRE in the 3’UTR of TLR4. To further demonstrate that TLR2 is responsible for the increased endothelial activation response, TLR2 receptors were blocked using TLR2 antibody and the effect on PBMC transmigration across the infected monolayer was analyzed. Blocking of TLR2 receptors resulted in a significant decrease in the percentage of transmigrating CD14^+^ cells across the DENV2-infected monolayer as compared to the controls (Figure 7g-7j). Therefore, blocking of TLR2 receptors in endothelial cells replicated the similar effect observed during miR-573 overexpression in DENV2-infected HUVEC.

**Figure 7:**
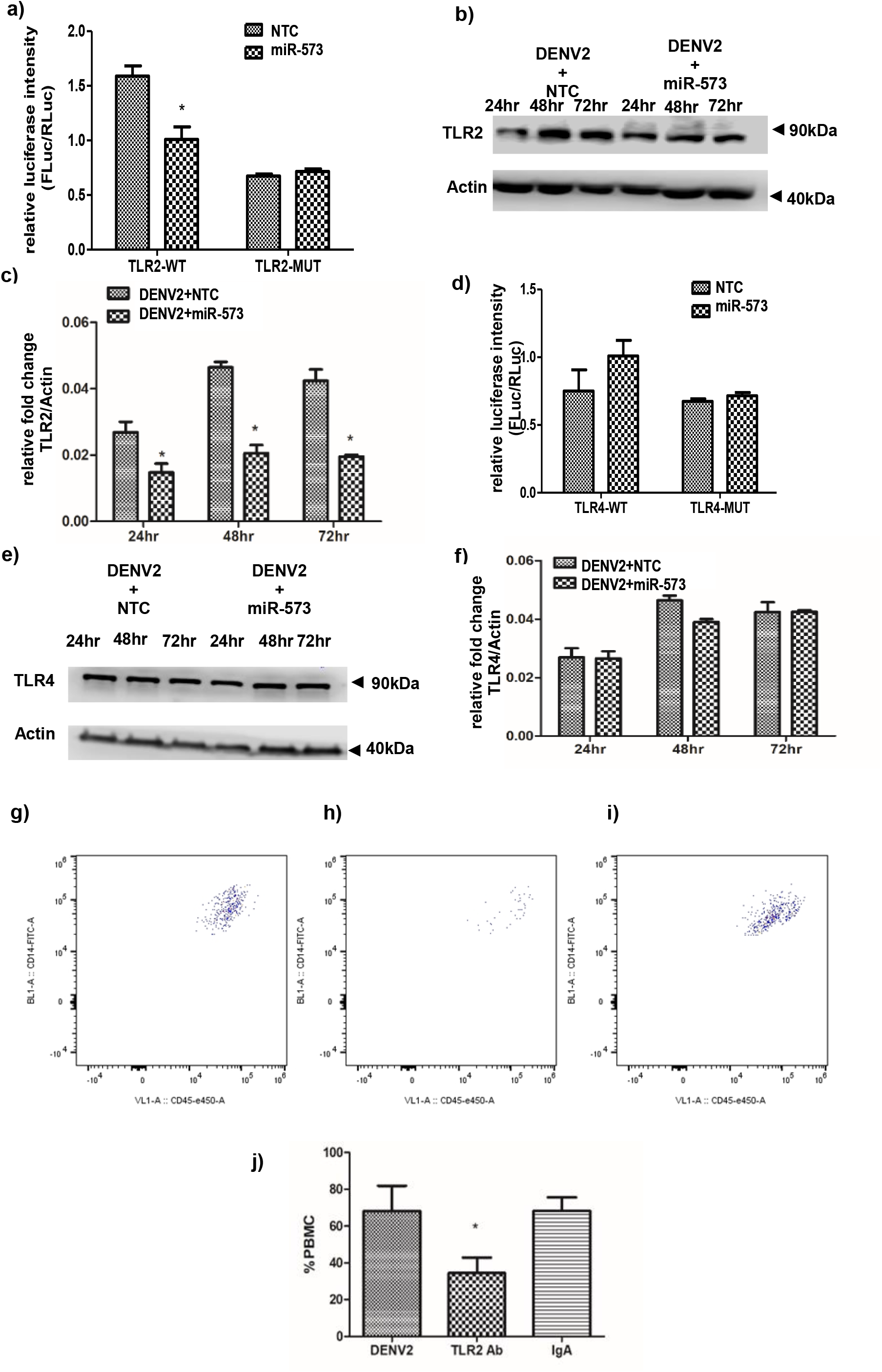
miR-573 targets TLR2 expression and modulates the endothelial activation response in DENV2-infected HUVEC. **a)** HUVEC cells were co-transfected with WT-3’UTR-TLR2 construct and miR-573 mimic (80nM) or equimolar concentration of non-targeting control (NTC) and the luciferase signal intensity was measured at 48 hpt. **b)** HUVEC cells transfected with miR-573 mimic or the NTC were infected with DENV2 at MOI 10 and cell lysates were collected at the indicated time intervals post infection for western blot analyses. **c)** A bar graph depicting the densitometric analysis of the decrease in TLR2 protein expression in miR-573 mimic transfected cells as compared to the NTC. **d)** Bar graph for luciferase expression from 3’UTR TLR4 constructs upon miR-573 mimic and NTC treatment (similar to TLR2). **e)** Western blot for TLR4 protein expression in DENV2-infected HUVEC cells upon miR-573 overexpression. **f)** A bar graph depicting the densitometric analysis of TLR4 protein expression. Blocking TLR2 decreases transendothelial migration of PBMCs across the DENV2-infected HUVEC monolayer. Representative images depicting the population of CD14 and CD45 positive cells transmigrated upon **(g)** DENV2 infection, **(h)** anti-TLR2 (1μg/ml) treatment and **(i)** IgG treatment (1μg/ml). **j)** A bar graph depicting the percentage of PBMC transmigrating across the infected HUVEC monolayer subjected to the different treatments. Data represented mean ± SEM and are obtained from three individual biological replicates. Student’s t test was carried out between individual groups to determine the statistical difference. *\*P<0*.*05*.

## DISCUSSION

Our study was the first to show that DENV2 infection in HUVEC results in differential expression of miRNAs which are potentially involved in regulating endothelial function and immune responses. We also showed that miR-573 reduces endothelial permeability by suppressing the expression of vasoactive ANGPT2 expression during DENV2 infection. It further also suppresses the exaggerated proinflammatory IFN response by downregulating TLR2 expression in DENV2-infected endothelial cells. We also identified that miR-573 was under the direct transcriptional control of nuclear hormone receptor PPARγ which plays an important role in maintaining endothelial cell function. This study identified a novel mechanism through which miR-573 and PPARγ rescue the endothelial dysfunction observed during DENV2 infection.

Earlier in vitro studies in non-endothelial cell types as well as in dengue patients have established that DENV infection modulates miRNA expression. However, none of the previous work have looked at the changes in miRNA profiles with respect to endothelial cells. Our study revealed that several miRNAs were down-regulated in HUVEC upon DENV2 infection. Earlier studies in Huh7 cells have shown that DENV NS3 and NS4B proteins were responsible for downregulating the major components of the host RNAi machinery which include the Dicer, Drosha, DGCR8 and Ago proteins [50]. Such a shutdown of the RNAi machinery would suppress miRNA biogenesis observed in DENV2-infected HUVEC as well. However, we did not study the effect of these DENV non-structural proteins on RNAi machinery and needs to be further examined whether similar mechanisms are in play in DENV2-infected endothelial cells. Our RNA-seq analysis also identified some of the previously studied miRNAs like miR-133a and miR-548g-3p which directly bind to the 3’ and 5’UTR of DENV genome and regulate virus replication [51, 52]. miR-21, miR-30e*, miR-378, miR-146a, miR-223 and let-7 family of miRNAs have been shown to indirectly regulate viral replication by targeting host factors involved in proinflammatory and innate immune responses [24-26, 51, 53-55]. miR-146a, which was also upregulated in DENV2-infected HUVEC, targets TRAF6 expression subverting the IFNβ and autophagy responses and enhances DENV replication in monocytes [25, 53]. let-7c and miR-21 were also found to be elevated in HUVEC with probable promote DENV replication as previously explored [26, 54]. miR-150 which suppresses SOCS1 expression and promotes proinflammatory pathogenic response during DENV infection was also found to be elevated in HUVEC cells upon DENV2 infection [56]. In addition, we also identified miRNAs (miR-26, miR-30, miR-130, miR-365, and miR-483) associated with endothelial function which were differentially regulated in DENV2-infected HUVEC. Interestingly these miRNAs were significantly downregulated at 24 hpi as compared to the other time points during infection. We hypothesize that the reduction in miRNA expression may correlate with the increased viral replication at 24 hpi (data not shown). These observations suggest that certain common mechanisms of miRNA mediated regulation of immune responses might also be at play in both endothelial and non-endothelial cell types during DENV infection.

We further focused on a miR-573 which was identified as a potentially important miRNA targeting endothelial functional responses in our gene ontology analysis. miR-573 has previously been explored in the context of different types of cancers with either tumorigenic or antimetastatic function [57-59]. miR-573 was also shown to suppress the proinflammatory response in synovial fibroblast cells of rheumatoid arthritis (RASF) thioredoxin domain containing 5 (TXNDC5). Addition of conditioned medium from miR-573 overexpressing RASFs decreases the angiogenic capacity of HUVEC [36]. We observed that DENV infection resulted in a decrease in miR-573 expression in HUVEC which did not show any dependence on the MOI. Infection with a replication defective UV-inactivated virus also resulted in the downregulation of miR-573 expression which further suggests that its expression does not depend on active viral replication. We observed that transient overexpression of the different DENV structural proteins resulted in the downregulation of miR-573 expression and the viral capsid protein had the most significant effect. However, we did not observe a dose-dependent response in miR-573 expression in response to the viral capsid protein expression and the other structural proteins (Figure 2 – figure supplement 1). We postulate that the presence of the DENV2 structural proteins in the cell trigger certain host intrinsic factors which might be a limiting factor in regulating miR-573 expression [60]. In fact our group has previously shown that DENV2 capsid protein alters HMGB-1 translocation in monocytes initiating a proinflammatory response [61]. Though we proposed that miR-573 expression does not depend on active virus replication because both wild type and UV-inactivated viruss resulted in decrease in miR-573 expression in comparison to mock-infected cells, the observed difference may have resulted from other factors released from infected cells into the medium. To further confirm that miR-573 expression in replication independent experiments with DENV2 VLPs (virus like particles) are necessary.

miR-573 is an intergenic miRNA which is located between DHX15 and PPARGCA1 on chromosome 4 with its own transcription regulatory elements. We identified a peroxisome proliferator-activated receptor gamma (PPARγ) response element (PRE) upstream of the transcription start signal (TSS) in the miR-573 coding region. PPARγ has an endothelial protective function and prevents vascular injury during proinflammatory vascular diseases like sepsis [62, 63] but not much is known about its role during DENV2 infection.. DENV2 infection resulted in a decline in PPARγ mRNA and protein levels and PPARγ activity positively correlated with miR-573 expression in DENV2-infected HUVEC. Treating HUVEC cells with a PPARγ agonists prior to DENV infection resulted in greater increase in miR-573 transcript levels as compared to treating the cells post infection. No change in miR-573 transcript levels was observed in DENV2-infected cells when treated with PPARγ antagonist (Figure 3 – figure supplement 2) as opposed to the decline observed in mock-infected cells. The lack of significant decrease in miR-573 expression due to PPARγ antagonism may be attributed to the already low expression of PPARγ and miR-573 in DENV2-infected HUVEC. We further confirmed that transient over-expression of PPARγ in DENV2-infected HUVEC increased miR-573 expression. Furthermore, PPARγ was demonstrated to bind upstream to the miR-573 coding region by chromatin immunoprecipitation assay. Therefore, these observations suggest that PPARγ positively regulates miR-573 expression at the transcriptional level and the decrease in PPARγ expression in DENV2-infected HUVEC is probably responsible for downregulation of miR-573. It has previously been shown that PPARγ expression is affected by the oxidative stress response in endothelial cells induced by proinflammatory stimuli. PPARγ mRNA transcription and activity were inhibited in HUVEC during H_2_O_2_ exposure which may be mediated by redox dependent transcription factors like AP1 [38]. DENV2 infection in HUVEC is also associated with an increase in ROS levels which play a role in endothelial dysfunction [64]. We observed that inhibiting ROS production in DENV2-infected HUVEC cells increases PPARγ and miR-573 expression. Therefore, the ROS imbalance induced during DENV2 infection in HUVEC, decreases PPARγ expression resulting in reduced transcriptional activation of miR-573 (Figure 8a). However, we observed that at 48 hpi miR-573 expression was reduced whereas total PPARγ expression remained high in NAC treated cells. The decrease in miR-573 expression may be attributed to lower levels of PPARγ nuclear translocation. The ROS levels in NAC treated cells at 48hpi are probably higher which might result in reduced PPARγ activity. We further showed that DENV2 structural proteins and particularly the capsid protein was responsible for the downregulation of miR-573 expression. We need further experiments to explore whether DENV2 capsid protein induces an oxidative stress response in DENV2-infected HUVEC and downregulates PPARγ and miR-573 expression.

**Figure 8:**
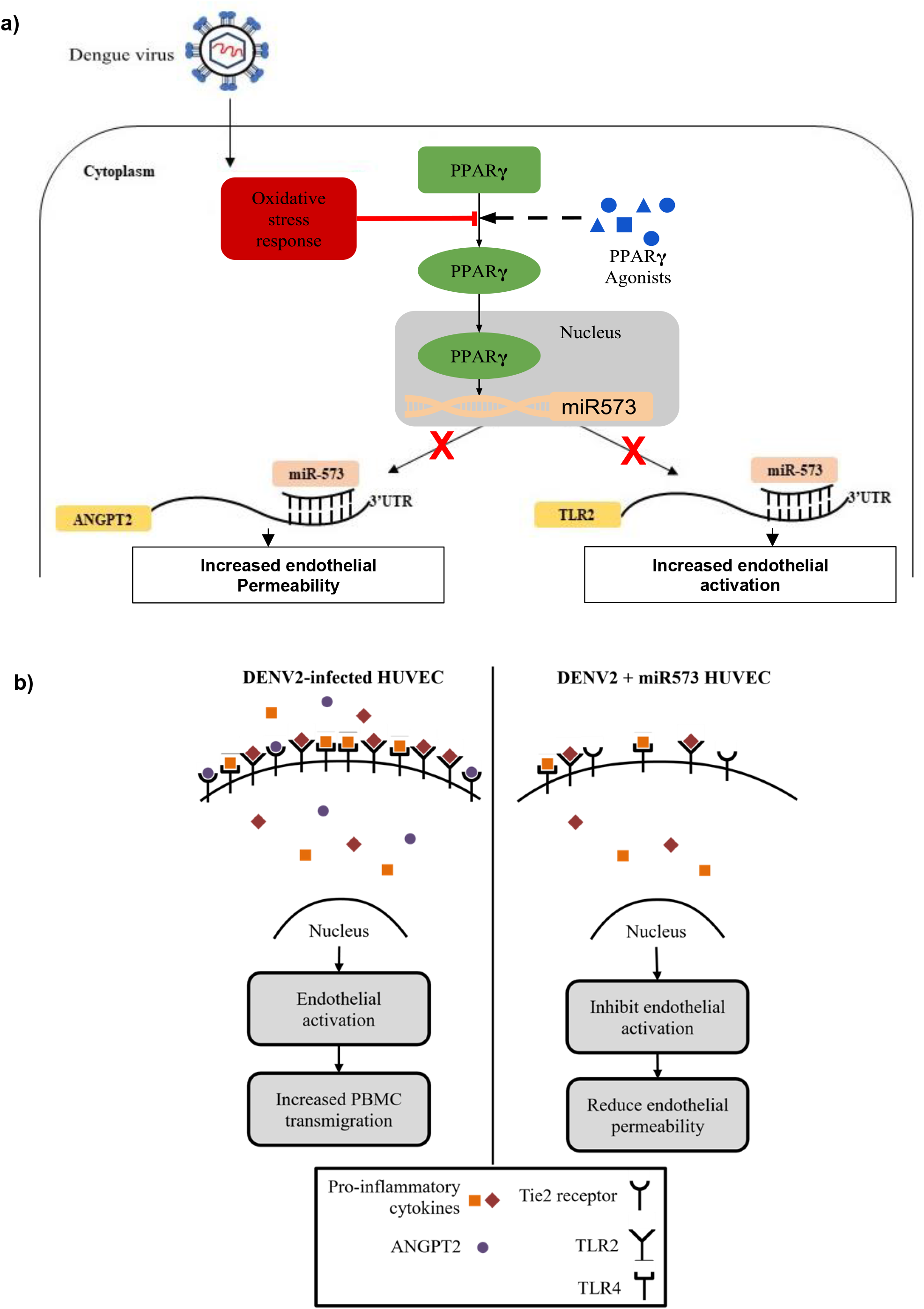
An overview of PPARγ mediated regulation of miR-573 and the downstream mechanism of regulating endothelial function DENV2 infection. Under normal homeostatic conditions PPARγ activation by endogenous agonists results in nuclear translocation of PPARγ where it binds upstream to the miR-573 coding region and drives its transcription. During DENV2 infection, the resulting oxidative stress response in endothelial cells inhibits PPARγ activation and nuclear translocation. As a result, PPARγ binding to the miR-573 promoter region is inhibited resulting in downregulation of miR-573 expression. During DENV2 infection, downregulation of miR-573 expression results in increased expression of its targets ANGPT2 and TLR2. ANGPT2 increases endothelial permeability by disrupting the adherens junction organisations in the endothelial cells. In addition, increased expression of TLR2 on the surface of the DENV2-infected cells results in increased interactions with vasoactive mediators, consequently mediating proinflammatory signalling through the TLR pathway. Overexpressing miR-573 in DENV2-infected HUVEC restores the functional activity of the miRNA and results in post transcriptional repression of ANGPT2 and TLR2 expression. miR-573 has an endothelial protective function and is involved in mitigating the activation and permeability response during inflammation.

We observed that overexpressing miR-573 in DENV2-infected HUVEC reduces endothelial permeability as observed with the increase in transendothelial electrical resistance and decrease in dextran FITC influx We also observed comparatively more intact VE-cadherin junction complexes as in miR-573 overexpressing DENV2-infected HUVEC as compared to the NTC transfected cells. However, immunofluorescence staining of the VE-cadherin complexes is only a qualitative assessment of their organization at inter-endothelial junctions and would further require higher resolution confocal microscopy to validate these findings. miR-573 had no effect on DENV2 viral replication, suggesting that the decrease in endothelial permeability is not due to decreased viral load and is probably mediated through host factors regulating endothelial function. ANGPT2 was identified as a predicted target of miR-573. Angiopoietins (ANGPT1 and ANGPT2) are endothelial growth factors which regulate endothelial cell homeostasis through Tie receptor tyrosine kinases [45, 65]. ANGPT1 is associated with maintaining endothelial cell stability, integrity and inducing an anti-inflammatory response [45]. In contrast, ANGPT2 antagonizes the protective effect of ANGPT1 signaling by engaging the common Tie2 receptor. Under normal cellular conditions ANGPT1 levels are higher than ANGPT2 and an imbalance in ANGPT1/ANGPT2 ratio is associated with endothelial dysfunction in vascular inflammatory diseases like sepsis [45]. In vitro as well as clinical studies have shown that during DENV2 infection ANGPT2 levels are elevated resulting in a skewed ANGPT1/ANGPT2 ratio [30, 66]. Increase in ANGPT2 expression is associated with disorganized VE-cadherin complexes at intercellular junctions in the endothelium [65]. Additionally, under proinflammatory conditions ANGPT2 induces the formation of actin stress fibers resulting EC contractility and destabilizes EC-ECM adhesion through α5β1 integrin signaling [67]. The factors responsible for the elevated ANGPT2 expression are less explored. miR-573 was predicted to bind to the miRNA response element (MRE) present in the 3’UTR of ANGPT2. Additionally, the expression levels of miR-573 and ANGPT2 were negatively correlated. miR-573 was shown to bind to the 3’UTR of ANGPT2 in the luciferase reporter construct which resulted in decreased luciferase expression. Overexpressing miR-573 suppressed ANGPT2 expression and reduced the secreted levels of ANGPT2 from DENV2-infected HUVEC. Furthermore, addition of recombinant ANGPT2 protein to the DENV2-infected cells abrogated the protective effect of miR-573 resulting in increased endothelial permeability. This suggests that the decrease in endothelial permeability mediated by miR-573 was due to the post transcriptional repression of ANGPT2 protein expression and addition of recombinant ANGPT2 protein overcame this repressive effect in the infected cells.

miR-573 was also responsible in mitigating the endothelial activation response in DENV2-infected HUVEC. Activated endothelial cells promote leukocyte transcytosis which further contribute to the local proinflammatory response thus, exacerbating endothelial permeability. Over-expressing miR-573 resulted in decreased expression of proinflammatory cytokines like IL-6 and IFN response genes which promote monocyte recruitment and platelet attachment to the activated endothelium. Increase in circulating levels of IL-6 in patient sera strongly correlates with increased propensity for developing vascular leakage [68]. A reduction in endothelial activation also resulted in a decrease in the percentage of PBMC transmigrating across the DENV2-infected endothelium. Several proinflammatory factors, which are induced by different innate immune mechanisms, increase endothelial activation response in DENV2-infected HUVEC. TLR signaling is an innate immune response mechanism which has been linked to endothelial activation and dysfunction [69-71]. TLRs (TLR2, TLR4 and TLR6) are required for DENV2 NS1 induced endothelial activation response [49, 72]. Other vasoactive factors like HMGB1, IL-1β, TNFα, histamine etc., which are significantly elevated during DENV infection, also mediate endothelial permeability via TLR signaling pathways [72, 73]. miR-573 was predicted to target the 3’UTR of TLR2 and TLR4 which were over-expressed in DENV2-infected HUVEC. In vitro studies (as previously described) showed that miR-573 directly binds to the 3’UTR of TLR2 and decreases its expression at both mRNA and protein level. Though miR-573 resulted in a decrease in luciferase expression in the TLR4 3’UTR luciferase construct, overexpressing miR-573 did not result in a decrease in TLR4 mRNA and protein expression. Furthermore, blocking of TLR2 in DENV2-infected endothelial cells replicated the effect of miR-573 on Il-6 secretion and PBMC transendothelial migration. These observations suggest that miR-573 suppresses TLR2 expression thus reducing its surface expression. Lack of availability of TLR2 receptors on the endothelial cell surface regulates the engagement of these limited receptors with vasoactive factors limiting the endothelial activation response.

The current study shows that miR-573 plays a dual role of reducing the endothelial permeability and activation responses during DENV2 infection (Figure 8b). It would be interesting to further explore whether miR-573 exhibits similar endothelial protective function in response to other proinflammatory stimuli as well as DENV NS1 treatment. Further in vivo and clinical studies are warranted for determining the therapeutic potential of miR-573 in rescuing vascular leakage during severe dengue infection. However, absence of a miR-573 homologue in mice hampers further in vivo studies. An indirect approach of modulating the PPARγ activity to reduce vascular leakage in dengue mouse models can also be explored. PPARγ agonists are already being used as insulin sensitizers in diabetic patients and their clinical potential is further being explored with respect to other metabolic and inflammatory diseases [74, 75]. It would be interesting to explore whether PPARγ agonists can also be repurposed as therapeutic agents to prevent endothelial permeability in dengue patients at the risk of developing vascular leakage.

## Key Resources Table

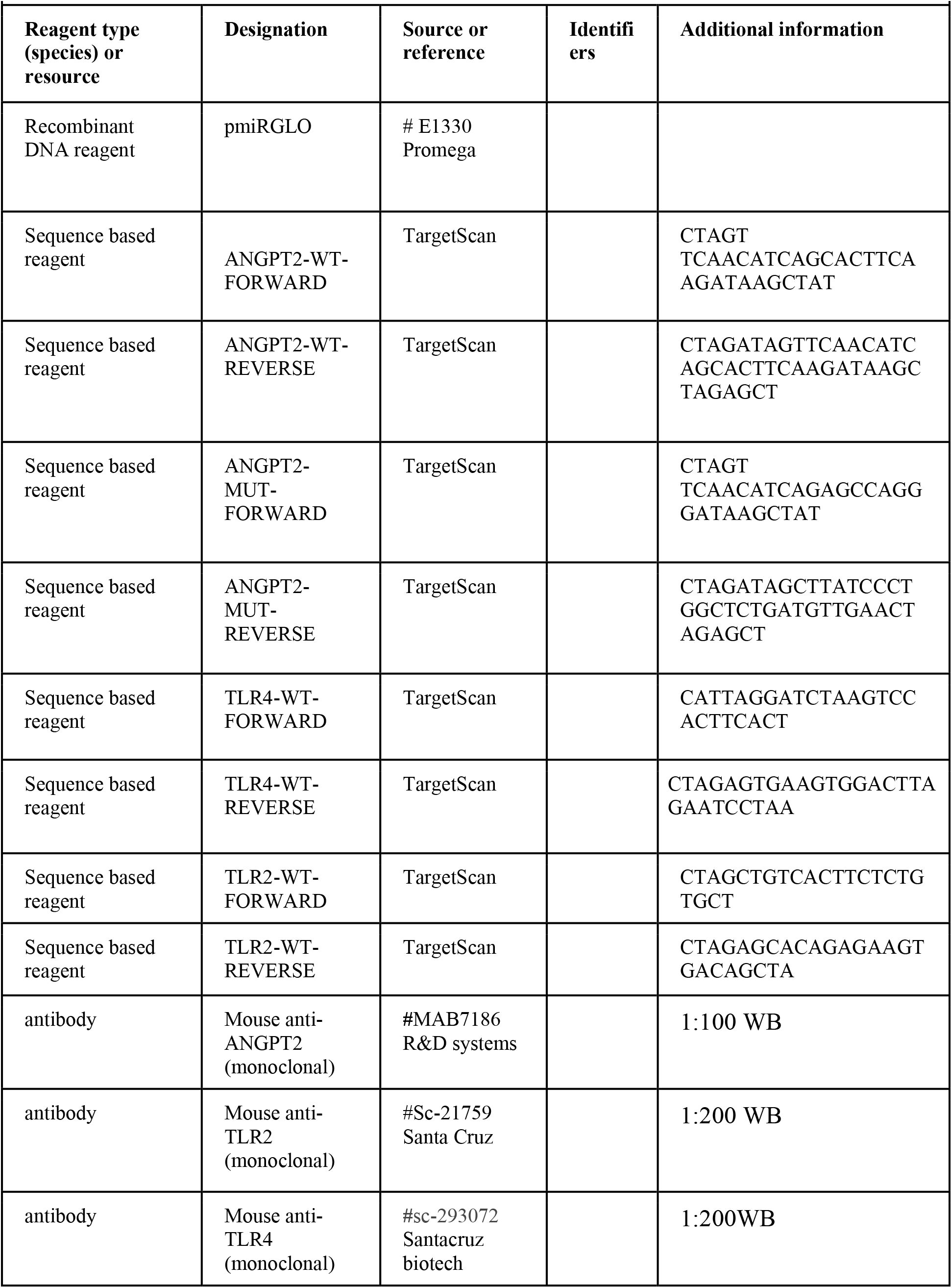
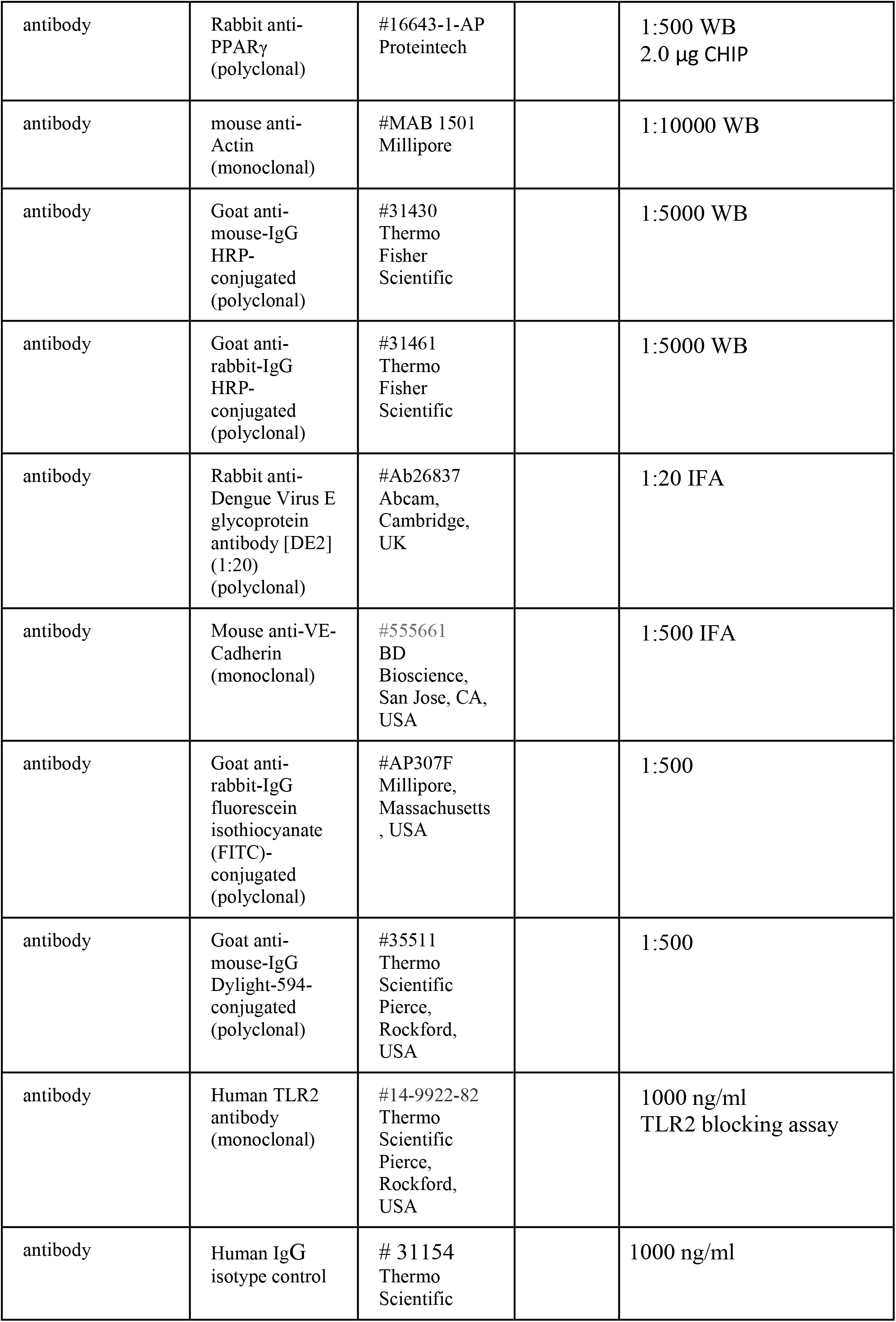
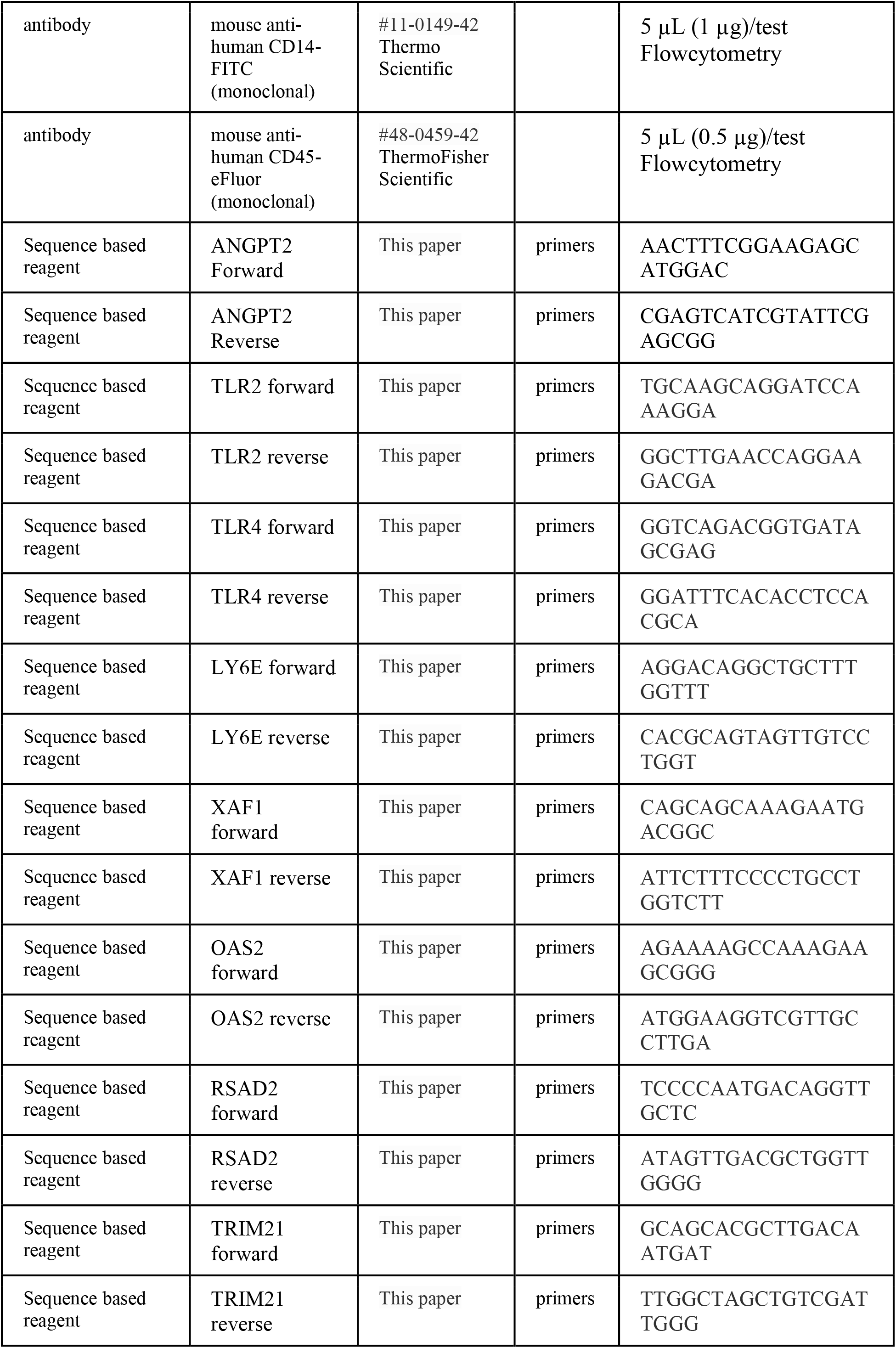
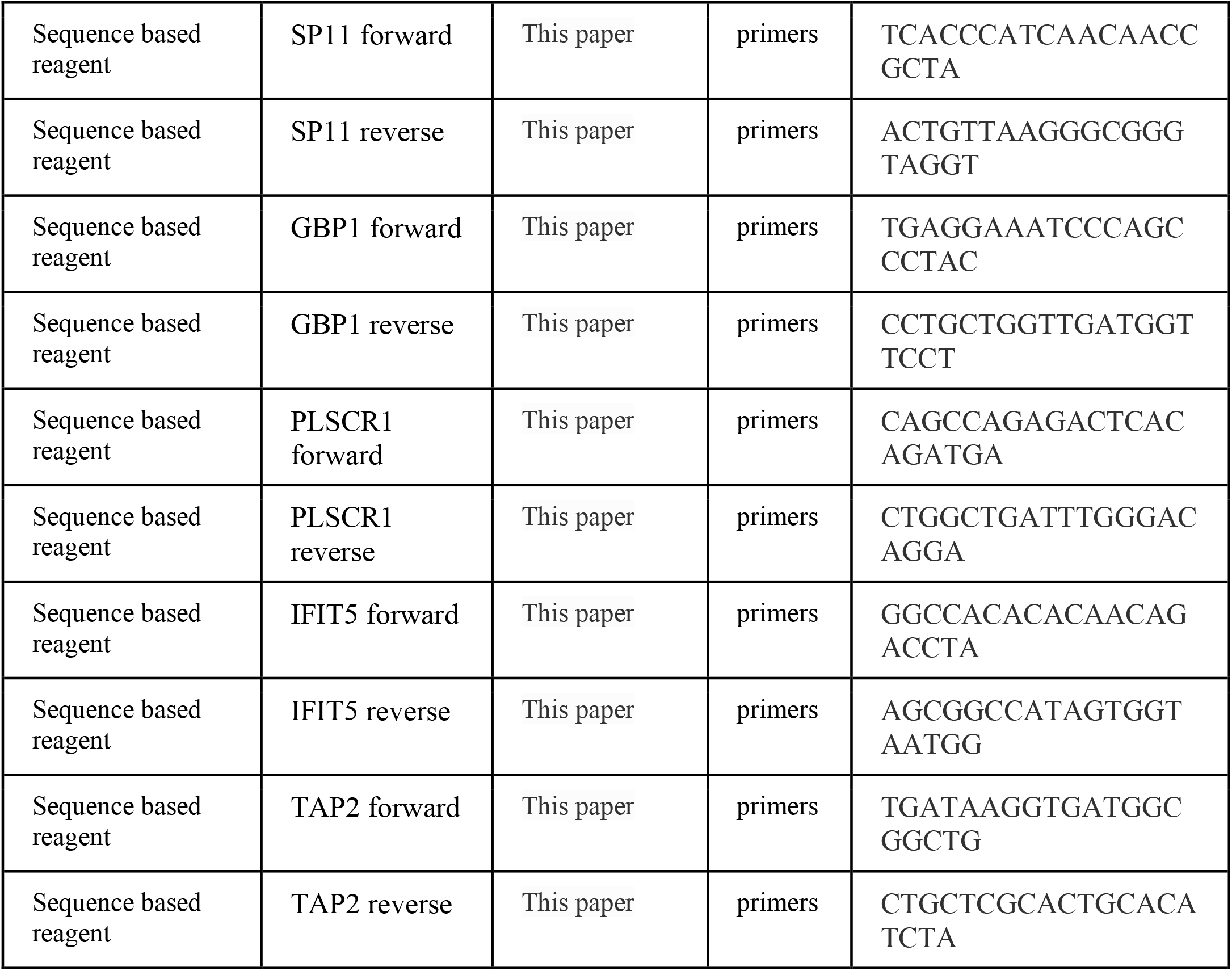

## Material and Methods

### Virus propagation and in vitro infection

DENV2 (strain Den2STp7c6) was propagated in C6/36 cells and viral titers were quantified through plaque assays on BHK-21 cells.

HUVEC (C2517A, Lonza) and HMEC-1(CRL-3243, ATCC) cells were seeded at 5 × 10^4^ cells/cm^2^ density and upon reaching 80% confluency were infected with DENV2at the desired MOI and incubated at 37°C for 1 hr. The cells were then washed with sterile PBS and overlaid with EBM-2 media supplemented with 10% FCS and incubated at 37°C with 5% CO_2_. At the desired time intervals supernatants were collected and the virus titers were quantified using plaque assays as previously described. All the cells used in this study tested negative for mycoplasma contamination using MycoAlert™ Mycoplasma detection kit (Lonza).

### Next Generation miRNA sequencing and microarray analysis

Total RNA from DENV2-infected and mock-infected cells were isolated using mirVANA RNA isolation kit (Thermofisher). The Illumina HiSeq2500 platform was used for miRNA sequencing. Briefly, cDNA libraries for the RNA samples were generated using the TruSeq Small RNA library preparation kit (Illumina). The cDNA libraries size selected (125 – 175 bp) and mixed in equimolar concentrations and sequenced using the standard TruSeq primer with 50 bp paired end reads. The processed reads were analysed using for differential miRNA expression, miRNA lists were created using a cut-off of P<0.05, >/<2-fold change. Illumina microarray using the Human HT-12 v4.0 BeadChip platform was used for differential gene expression studies. Gene expression data were analysed using DESeq package (R Bioconductor 3.9) and gene lists were created using the cut-off of P<0.05, >/< 2-fold change. All the raw and processed data files are deposited in the NCBI GEO database (GSE135311).

### Endothelial permeability and leukocyte transmigration assays using transwell inserts

HUVEC seeded in transwell inserts (0.3 µm pore, Thincert™ Grenier) were infected with DENV at MOI 10 upon reaching confluency. Paracellular permeability was assessed by the addition of 0.5 mg/ml fluorescein isothiocyanate-conjugated 70-kDa dextran (dextran FITC, Sigma) to the upper chamber and incubated at 37°C for 30 mins. 100 µL of medium was removed from the lower chamber for fluorometric analysis 490 nm/520 nm. The transendothelial electrical resistance (TEER) of the HUVEC monolayers was measured with Millicell-ERS volt-ohmmeter (Millipore) and reported as percentage difference between TEER values before and after infection/treatment.

Leukocyte transmigration assay was carried out by addition of 5×10^6^ cells/mL hPBMC (#CC-2702 Lonza) to DENV2-infected HUVEC monolayers at 24 hpi following different treatments. For blocking of TLR2 receptors, DENV2-infected HUVEC were incubated with hTLR2 and IgG isotype control for 30 mins prior to PBMC addition. After 3 hr of incubation at 37°C, media containing the transmigrated PBMC was collected from the lower chamber and the cells were fluorescently labelled CD14-FITC and mouse anti-human CD45-eFluor for flowcytometric analysis.

### miRNA LNA-mimic, LNA-inhibitor and plasmid transfection

HUVEC cells were reverse transfected with 80 nM of miR-573 mimic or equimolar concentration of cel-miR-39-3p, a mimic negative control (Qiagen) using 4 µM/mL Endoporter reagent (GeneTools,LLC). DENV2 structural proteins were cloned into expression vector from Promega in frame with a C-terminal Flag tag. PPARγ cDNA clone with the C-terminal Flag tag was purchased from (HG12019-CF, Sino biological). The 3’UTR luciferase constructs were designed using the pmiRGLO Dual luciferase miRNA target expression vector. The wild-type and mutant miRNA target sites were cloned downstream of the Firefly luciferase (luc2) reporter gene in pmirGLO vector (Promega) for luciferase reporter assays. The plasmid constructs were transfected using the Amaxa™ 4D-Nucleofector™ CA-167 (Lonza) protocol.

### Real-time-polymerase chain reaction qRT-PCR Analysis

Total RNA isolated from HUVEC was used for miRNA quantification using the miRCURY™ LNA™ miRNA RT Kit (Qiagen). Briefly, cDNA synthesis was carried out for 60 min at 42°C followed by a heat-inactivation step at 95°C for 5 min. The cDNA was diluted with RNase-free water (1:60) and used for PCR reaction. The cycling conditions used are as follows, heat activation for 2 min at 95°C followed by denaturation for 10 sec at 95°C, combined annealing and extension at 56°C for 60 sec for 40 cycles. The melt curve analysis was carried out between 60-95°C. For gene expression analysis one-step qRT-PCR (SYBR Green Quantitative RT-PCR kit, Sigma) was carried out. cDNA synthesis was carried out at 42°C for 30 min followed by denaturation and heat inactivation of RT enzyme at 95°C for 30 sec. The PCR cycling conditions used for amplification are as follows, denaturation at 95°C for 5 secs, primer annealing at 55°C for 15 sec and extension at 72°C for 10 sec and were carried out for 40 cycles. The melt curve analysis was carried out between 60-95°C. Data analysis was carried out using the software supplied with Applied Biosystems StepOne Plus.

### Protein estimation and indirect immunofluorescence assay

For subcellular fractionation studies cell-lysis buffer was added to HUVEC cells and incubated for 15 min on ice to allow for complete lysis. The cell lysates were centrifuged at 720g for 5 min at 4°C to separate the nuclear and cytoplasmic fractions. The nuclear pellet was washed twice with 500 µL lysis buffer followed by centrifugation at 720xg for 10 min. The pellet was resuspended in Tris buffered saline (TBS) containing 0.1% SDS and protease inhibitor before proceeding to sonication. Protein estimation of the nuclear and cytoplasmic fractions was carried out by Bradford assay and then used for western blot analysis. Alternatively, 1% SDS cell-lysis buffer was used for cell lysis and protein extraction for western blot analysis. Densitometric analysis was carried out using the C-digit western blot scanner (LI-COR) and the relative protein expression was normalized to actin loading controls. The amount of ANGPT2 secreted from HUVEC into the cell culture supernatant was quantified using Human Angiopoietin 2 DuoSet ELISA kits (R&D systems), following the manufacturer’s instructions. DENV2-infected HUVEC were fixed and permeabilized using 4 % paraformaldehyde (PFA) and 0.01% TritonX-100 for 15 min and then washed three times with PBS.

### Chromatin Immunoprecipitation (ChIP) assay

HUVEC cells were crosslinked with 0.75% formaldehyde for10 min at room temperature with constant shaking followed by 0.125 M glycine addition to terminate crosslinking. Fixed cells were washed with ice-cold PBS and centrifuged at 1000xg for 5 min at 4°C. The pellet is resuspended in 500 µl ChiP lysis buffer and sonicated to shear DNA to 250–1000 base-pairs. The cell debris was pelleted by centrifugation for 10 min, 4°C, 8,000xg, and the supernatant containing the chromatin complex was incubated with the anti-PPARG antibody overnight at 4°C with continuous agitation. Protein G Magnetic beads (Millipore) were used for pull-down of PPARG-chromatin complex and analysed using PCR after reversal of cross-linking.

## Acknowledgements

This study was supported by Ministry of Education (MOE), Singapore Tier 2 (MOE-2017-358 T2-1-078) grant.

---

**Figure1 – supplement 1: DENV2 infection of endothelial cells results in differential expression of miRNAs**.Differentially expressed miRNAs in DENV2-infected HUVEC **a)** A venn diagram representing the miRNAs which were previously reported to be differentially regulated in other DENV2-infected non-endothelial cells. **b)** A volcano plot where the x-axis represents the logarithm to the base 2 of the fold changes and the y-axis specifies the logarithm to the base 10 of the t-test p values. The dots indicate miRNAs which passed the filtering criteria (log2 fold change +/- 2 and p-value <0.05). Validation of differentially expressed miRNAs by qRT-PCR. The expression profiles of up and down-regulated miRNAs obtained from RNAseq analysis were validated by qRT-PCR The **c)** up and **d)** downregulated miRNAs showed similar trends in expression for both RNAseq and qRT-PCR quantification methods. Putative pathways affected by the dysregulated miRNA response in ENV2-infected HUVEC. Diana mirPath pathway enrichment analysis was carried out to determine the biological pathways affected by the dysregulated miRNA expression during DENV2-infection, Fisher’s exact test (p < 0.05) and False discovery rate (FDR=0.01) **e)** Bar graph representing the distribution of miRNA target genes in different cellular pathways. f) qRT-PCR analysis of miR-573 transcript levels in HMEC-1 cells when infected with DENV2 at different time intervals post infection.

**Figure 2 – supplement 1: DENV2 structural proteins downregulate miR-573 expression**. HUVEC cells were transiently transfected with recombinant DENV2 **a)** envelope **b)** capsid and **c)** prM gene constructs and miR-573 expression was quantified using qRT-PCR. Western blot analysis was carried out to confirm the expression of each of the viral proteins. Data represented are obtained from biological triplicate experiments Student’s t test was applied to determine the statistical difference between individual groups \*\**\*P<0*.*001*.

**Figure 3 – supplement 1**: **Computational analysis of PPARγ binding site upstream of miR-573 coding region. a)** CAGE tags, TSS Seq tags, H3K4me3 modification, ESTs and conservation patterns are clearly displayed in the 50 kb upstream region of hsa-miR573. The red line in the block of putative TSS denotes the representative TSS, whereas the blue lines denote other TSS candidates of hsa-miR573. **b)** The predicted PPARγ transcription response element is located 1536 bp upstream of the miR573 gene locus **c)** Diagrammatic representation of the regulatory interaction between PPARγ and miR573 at the genetic level.

**Figure 3 – supplement 2: PPARγ expression upon DENV2 infection and NAC treatment**. HUVEC cells were treated with 10 µM of rosiglitazone **a)** pre and **b)** post DENV2 infection at MOI 10 and miR-573 expression was measured using qRT-PCR. **c)** miR-573 expression was also quantified in DENV2-infected cells treated with 10 µM of GW9662 prior to DENV2 infection. No significant decrease in miR573 transcript levels was observed for between cells treated with 10 µM of GW9662 and 0.01% DMSO (vehicle control) 24 hpi. HUVEC cells were treated with 5mg/ml of N-acetyl cysteine (NAC) 12 hrs prior to infection and then infected with DENV2 at MOI 10. The cells were constantly maintained in media containing 5mg/ml NAC. **d)** Addition of NAC resulted in a decrease in reactive oxygen species (ROS) in DENV2-infected cells when compared to the untreated cells. **e)** Cell lysates were analyzed for PPARγ protein expression at 24, 48 and 72 hpi by Western blot. **f)** A histogram depicting PPARγ protein levels were up-regulated in DENV2-infected cells treated with NAC.

**Figure 4 – supplement 1: Regulation of endothelial permeability response by modulating miR-573 expression**. HMEC-1 were transfected with miR-573 mimic (80nM) or the mimic non-targeting control (NTC, 80nM) were seeded onto the transwell inserts and endothelial permeability was measured following mock or DENV2 infection. **a)** TEER measurements across DENV2-infected endothelial monolayer transfected with miR-573 mimic increased gradually. **b)** Similarly, the dextran FITC influx across the DENV2-infected endothelial monolayer transfected with miR-573 mimic decreased. HUVEC cells were transfected with miR573 inhibitor (100nM) or the non-targeting control (NTC,100nM) and seeded on transwell inserts. **c)** TEER values were measured after 24 hpi with DENV2. Inhibition of miR573 activity in mock-infected HUVEC resulted in a significant decrease in TEER values by 26.5% from 24 to 48 hpi (n=3, *p<0.05, **p<0.01, **p<0.001). No significant decrease in TEER values was observed between DENV2-infected HUVEC treated with miR573 inhibitor or the NTC. Data represented are obtained from biological triplicate experiments. Student’s t test was applied to determine the statistical difference between individual groups *\*P<0*.*05, **P<0*.*01*

**Figure 5 – supplement 1: Increase in ANGPT2 expression during DENV2 infection causes vascular leak in endothelial cells and inhibition of miR-573 has no effect on its expression**. HUVEC cells were infected with DENV2 at MOI 10 and ANGPT2 transcript levels were measured by qRT-PCR at regular time intervals post infection. ANGPT2 mRNA transcripts were elevated by 13 and 11 folds elevated at 24 and 48 hpi respectively (n=5, ***p<0.001). **b)** ANGPT2 protein expression was also higher in DENV2-infected HUVEC at 24 and 48 hpi as compared to mock-infected cells. **c)** A bar graph showing the densitometric analysis of the western blot for ANGPT2 expression. DENV2 infection resulted in a 3.5 and 3.4 fold increase in ANGPT2 protein levels in HUVEC in comparison to mock-infected cells (n=3, *p<0.05). **d)** Secreted ANGPT2 levels were determined by ELISA which showed an increase in ANGPT2 secretion from DENV2-infected HUVEC (124.3 to 206 pg/ml) when compared to mock-infected cells (100.3 and 87.3 pg/ml, n=3, *p<0.05)). Addition of recombinant ANGPT2 (30 ng/ml) to DENV2-infected monolayer resulted in an increase in endothelial permeability as seen by **e)** the decrease in TEER values and **f)** increase in the dextran FITC influx. **g)** HUVEC cells were co-transfected with WT-3’UTR-ANGPT2 construct and miR573 inhibitor (100nM) or the non-targeting control (NTC,100nM) and the relative luciferase signal intensity was measured 24 hpt. The luciferase expression was 1.2 folds higher in the cells transfected with the inhibitor as compared to the non-targeting control. No significant difference was observed in the luciferase expression in between the inhibitor treated and the NTC treated cells co-transfected with the MUT-ANGPT2 construct. **h)** No significant difference was observed in the ANGPT2 protein expression between cells transfected with miR573 inhibitor and the NTC. Student’s t test was carried out to determine the statistical differences between individual groups. Data represented are obtained from biological triplicate experiments Student’s t test was applied to determine the statistical difference between individual groups *\*P<0*.*05, **P<0*.*01*.

**Figure 6-figure supplement 1: Regulation of proinflammatory response upon miR-573 inhibition**. HUVEC cells were transfected with miR573 inhibitor (100nM) or the non-targeting control (NTC,100nM). Inhibition of miR573 activity had no significant effect on **a)** IL-6 secretion **b)** and PBMC transmigration in DENV2-infected HUVEC as compared to the NTC. Data represented are obtained from biological triplicate experiments Student’s t test was applied to determine the statistical difference between individual groups *\*P<0*.*05, **P<0*.*01*

**Figure 7 – supplement 1: TLR2 expression is elevated during DENV2 infection in endothelial cells and inhibition of miR-573 activity has no effect on its expression. a)** TLR2 transcript levels transcript levels increased from 0.8 folds to 6.9 folds at 24 and 48 hpi respectively (n=3, *p<0.05). **b)** Western blot analysis showing the increase in TLR2 expression in DENV2-infected HUVEC. **c)** Densitogram analysis (bar graph) shows that TLR2 protein levels increased from 0.6 to 1.17 folds at 24 and 48 hpi respectively (n=3, *p<0.05). **d)** TLR4 mRNA transcript levels marginally increased from 1 to 1.08 folds at 24 and 48 hpi respectively (n=3, p<0.05). **e)** TLR4 protein expression was 3.1 and 2.1 folds higher at 24 and 48 hpi respectively in the DENV2-infected cells (n=3, p<0.05). **f)** Densitogram analysis (bar graph) highlighting the increase in TLR4 protein expression in DENV2-infected HUVEC. HUVEC cells were co-transfected with WT-3’UTR-TLR2 construct and miR573 inhibitor (100nM) or the non-targeting control (NTC,100nM) and the relative luciferase signal intensity was measured 24 hpt. **g)** Addition of miR573 inhibitor had no significant effect on the luciferase expression in cells transfected with WT-3’UTR-TLR2 construct as compared to the NTC. **h)** Inhibition of miR573 activity had no significant effect on TLR2 protein expression. Data represented are obtained from biological triplicate experiments. Student’s t test was applied to determine the statistical difference between individual groups *\*P<0*.*05*.

**Figure 4b source data file: Transendothelial resistance values (in ohms) after miR-573 mimic transfection in mock and DENV2-infected HUVEC**. The resistance values were recorded using Millicell-ERS volt-ohmmeter (Millipore) at the specific time points and each row represents an individual biological replicate.

**Figure 4c source data file: Dextran-FITC influx (Fluorescence units) after miR-573 mimic transfection in mock and DENV2-infected HUVEC**. At the specific time intervals (100 μL) of media from the lower chamber was collected and amount of dextran-FITC influx was measured through fluorometry. The excitation wavelength 490 nm and emission wavelength is 520 nm. Each row represents an individual biological replicate.

**Figure 5b source data file: Luciferase readings for ANGPT2 3’UTR constructs following miR-573 transfection**. Both Firefly and Renilla luciferase reading were recorded in pairs. Each row represents an individual biological replicate.

**Figure 5g source data file: Transendothelial resistance values recorded upon treatment with ANGPT2**. The resistance values were recorded using Millicell-ERS volt-ohmmeter (Millipore) at the specific time points and each column represents an individual biological replicate.

**Figure 7a source data file: Luciferase readings for TLR2 3’UTR constructs following miR-573 transfection**. Both Firefly and Renilla luciferase reading were recorded in pairs. Each row represents an individual biological replicate.

**Figure 7d source data file: Luciferase readings for TLR4 3’UTR constructs following miR-573 transfection**. Both Firefly and Renilla luciferase reading were recorded in pairs. Each row represents an individual biological replicate.

